# A novel small molecule that induces cytotoxicity in lung cancer cells inhibits disulfide reductases GSR and TXNRD1

**DOI:** 10.1101/2021.06.28.450088

**Authors:** Fraser D. Johnson, John Ferrarone, Alvin Liu, Christina Brandstädter, Ravi Munuganti, Dylan Farnsworth, Daniel Lu, Jennifer Luu, Tianna Sihota, Sophie Jansen, Amy Nagelberg, Rocky Shi, Giovanni C. Forcina, Xu Zhang, Grace S. W. Cheng, Sandra E. Spencer Miko, Georgia de Rappard-Yuswack, Poul H. Sorensen, Scott J. Dixon, Udayan Guha, Katja Becker, Hakim Djaballah, Romel Somwar, Harold Varmus, Gregg B. Morin, William W. Lockwood

## Abstract

High-throughput phenotype-based screening of large libraries of novel compounds without known targets can identify small molecules that elicit a desired cellular response, but additional approaches are required to find and characterize their targets and mechanisms of action. Here we show that a compound termed lung cancer screen 3 (LCS3), previously selected for its ability to impair the growth of human lung adenocarcinoma (LUAD) cell lines, but not normal lung cells, induces oxidative stress and activates the NRF2 signaling pathway by generating reactive oxygen species (ROS) in sensitive LUAD cell lines. To identify the target that mediates this effect, we applied thermal proteome profiling (TPP) and uncovered the disulfide reductases GSR and TXNRD1 as LCS3 targets. Through enzymatic assays using purified protein, we confirmed that LCS3 inhibits disulfide reductase activity through a reversible, uncompetitive mechanism. Further, we demonstrate that LCS3-sensitive LUAD cells are correspondingly sensitive to the synergistic inhibition of glutathione and thioredoxin pathways. Lastly, a genome-wide CRISPR knockout screen identified the loss of NQO1 as a mechanism of LCS3 resistance. This work highlights the ability of TPP to uncover targets of small molecules identified by high-throughput screens and demonstrates the potential utility of inhibiting disulfide reductases as a therapeutic strategy for LUAD.

## Introduction

Lung cancer is the leading cause of cancer mortality worldwide, mainly due to high incidence and the lack of effective therapeutic strategies for patients with advanced disease (1, 2). Targeted therapies that inhibit mutated components of key cellular pathways such as EGFR have proven successful in LUAD treatment (3–5); however, most patients are not candidates for these therapies and those that are, inevitably develop resistance to treatment (6). While the current array of targeted cancer therapeutics is directed against the products of some mutated oncogenes and components of their signaling pathways, growth or survival of cancer cells may also be dependent on the function of other genes that would provide additional targets for new classes of anticancer agents. Recent studies suggest that cancer cells can become reliant on genes that are not essential in normal cells—for example, to compensate for transformation-induced stress. Therefore, targeting the products of those genes in the context of a malignant phenotype could produce synthetic lethality (7, 8).

High-throughput screening of large libraries of small molecules, using physiologically-relevant assays in cancer and normal cell lines (9), provides a starting point for identifying novel cancer cell dependencies. Somwar et al. conducted a screen of 189,290 small molecules for their ability to inhibit the growth of human lung adenocarcinoma (LUAD) cell lines and described four compounds known as Lung Cancer Screen (LCS) 1 to 4 with certain favorable characteristics, including low molecular weight, the capability to readily cross the cell membrane, and a confirmed ability to inhibit the growth of LUAD—but not normal lung cells— at a low half-maximal inhibitory concentration (IC_50_) (10).

Superoxide dismutase 1 (SOD1) was previously identified as the molecular target of LCS1, demonstrating the utility of this approach for defining new dependencies in LUAD cells (11). In the current study, we have set out to identify novel dependencies by further characterizing LCS3, the previously unpublished structure of which is presented in Figure 1a. We have approached this issue in two ways—by describing the phenotype it elicits in susceptible LUAD cells and by defining the proteins with which it physically interacts. To circumvent some of the technical challenges associated with mapping a large number of drug-protein interactions, we have successfully employed the recently developed method called thermal proteome profiling (TPP). TPP entails subjecting protein lysates, in the presence and absence of a small compound, to a temperature gradient to precipitate insoluble heat-denatured proteins. The soluble fraction of protein is then recovered and the abundance of protein is quantified by performing tandem mass spectrometry (12, 13). It is expected that proteins interacting with the compound of interest will be stabilized or destabilized, leading to a change in the temperature at which the target protein denatures and precipitates out of solution. In this way, TPP exploits the thermodynamic changes to protein stability that occur upon binding to a ligand or small molecule and can be used to identify even rare proteins that are thermally affected by such interactions.

**Fig. 1.**
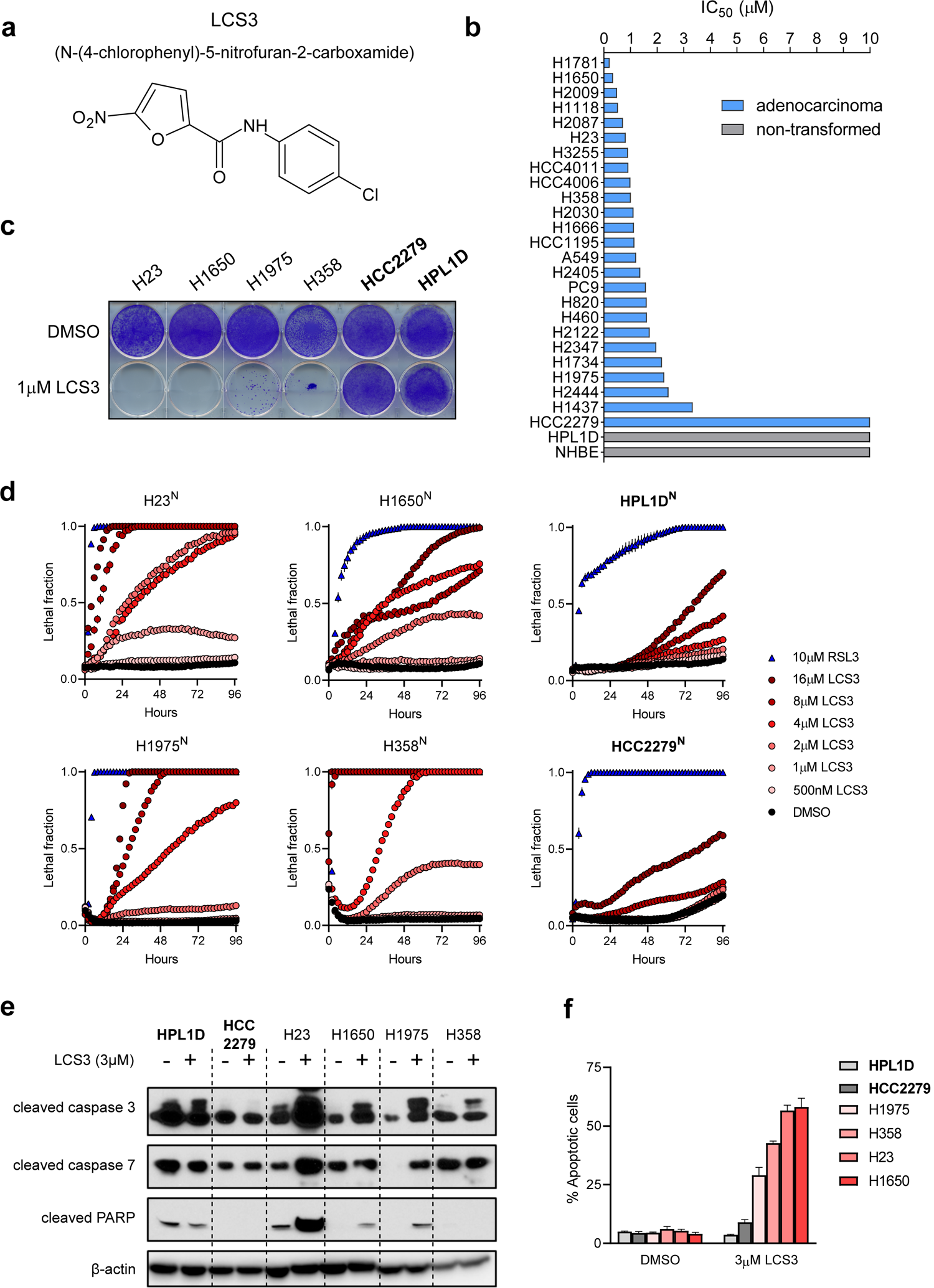
LCS3 inhibits lung cancer cell lines, but not non-transformed lung cells. **a-f** LCS3 resistant cell lines in bold. All error bars are +/- SEM. **a,** Molecular structure of LCS3. **b,** LCS3 sensitivity screen in LUAD and non-transformed lung cell lines. IC_50s_ determined by growth assays using alamarBlue cell viability reagent. **c,** LCS3 sensitivity determination by crystal violet staining to validate cell line screen results. **d,** Lethal fraction of lung cell lines treated with LCS3 determined by overlap of nuclei marker H2B-mCherry and dead cell marker SYTOX Green. **e,** Western blot of apoptotic proteins in lung cell lines treated with LCS3 for 96 hours. **f,** Apoptotic cell fraction of LCS3-treated lung cell lines determined by flow cytrometry detection of Annexin V stained cells.

Through the transcriptomic and proteomic profiling of gene expression, we found that sensitive LUAD cells accumulate ROS when exposed to LCS3, and we identified glutathione disulfide reductase (GSR) and thioredoxin reductase 1 (TXNRD1) as binding partners of LCS3. We then showed that LCS3 inhibits the enzymatic activities of both proteins *in vitro* and in cell-based assays. Because the combined inhibition of GSR and TXNRD1 exacerbates oxidative stress in LUAD cells and in non-malignant lung cells supplied with exogenous ROS, we suggest that high levels of ROS confer sensitivity to LCS3. Lastly, through a genome-wide CRISPR loss-of-function screen, we confirmed that the effects of LCS3 are related to antioxidant response and found that inactivation of NADPH quinone oxidoreductase 1 (NQO1) impairs ROS production and confers resistance to LCS3. Collectively, our results with LCS3 suggest that LUAD cells are reliant on the glutathione and thioredoxin reduction systems for viability and that targeting redox homeostasis may be a novel and effective therapeutic strategy in LUAD.

## Results

### LCS3 selectively inhibits LUAD cell viability

LCS3 (N-(4-chlorophenyl)-5-nitrofuran-2-carboxamide) (Pubchem ID: CID844044) is a heterocyclic compound (Fig. 1a) belonging to the 5-nitro-n-phenylfuran-2-carboxamide chemical class that was found to inhibit the proliferation of four LUAD cell lines, but not primary human bronchiolar epithelial cells (10). To assess the activity of LCS3 in a wider collection of LUAD cells, we screened a panel of twenty-seven cell lines for susceptibility to the effects of LCS3 on cell proliferation. Cells were exposed to concentrations of LCS3 ranging from 5nM to 10µM for 96 hours, and cell viability was determined using alamarBlue. Consistent with the original study (10), LCS3 selectively inhibited the growth of 26/27 LUAD cell lines at low micromolar concentrations (IC_50_ < 5µM); both of the non-transformed lung cell lines were relatively insensitive (IC_50_ > 10µM) (Fig. 1b).

To corroborate these results with an orthogonal approach, we performed clonogenic assays using six lung cell lines treated with 1 µM LCS3 for nine days. Consistent with the results shown in Figure 1b, we found that all LUAD cell lines except HCC2279 were sensitive to LCS3 while the non-transformed lung epithelial line HPL1D was resistant to treatment (Fig. 1c).

To determine the extent to which the loss of cell numbers is attributable to inhibition of proliferation or to induction of cell death, we used the scalable time-lapse analysis of cell death (STACK) method, which pairs live-cell imaging with fluorescence microscopy to detect the onset and rate of cell death (14). A fusion gene, H2B-mCherry, was transduced by lentivirus into lung cell lines to provide nuclear fluorescence; the resulting cell lines were denoted as H23^N^, H1650^N^, H1975^N^, H358^N^, HCC2279^N^, and HPL1D^N^. SYTOX Green—a nuclear dye which is unable to permeate the cell membrane of viable cells—was added to map the onset of toxicity in response to LCS3 or to a positive control lethal small molecule, the GPX4 inhibitor RSL3 (Supplementary Fig. 1a). We observed a more rapid increase in dead cells (the lethal fraction) in H23^N^, H1650^N^, H1975^N^ and H358^N^ cells than in HCC2279^N^ cells or the non-transformed HPL1D^N^ cells at most LCS3 concentrations, indicating that LCS3 inhibits cell growth through a mechanism that is cytotoxic, rather than cytostatic (Fig. 1d). We calculated the onset of cell death (D_o_) and the rate of cell death (D_R_) for each cell line; together, these data suggest that LCS3-sensitive cell lines undergo cell death that is both rapid in onset and synchronous throughout the cell population *in vitro* (Supplementary Fig. 1b).

We next sought to determine if the cytotoxicity induced by LCS3 occurs through an apoptotic mechanism. To this end, we treated cells with 3µM LCS3 for 96 hours and performed western blots for protein markers of apoptosis. We found that LCS3 increased cleavage of caspase 3, caspase 7 and/or PARP1 in all LCS3-sensitive LUAD cell lines (Fig. 1e). We also stained cells for annexin V and confirmed by flow cytometry that LCS3-sensitive, but not LCS3– resistant cell lines undergo apoptosis in response to 3µM LCS3 (Fig. 1f). Together, these data suggest that LCS3 selectively kills LUAD cell lines, at least in part through the induction of apoptosis.

To understand the ability of LCS3 to induce cell death across a breadth of cancer cell types, we determined the IC_50_ of LCS3 in the NCI-60 cell panel (Supplementary Fig. 1c). LCS3 caused toxicity across most types of cancer cell lines, with notable sensitivities in central nervous system and colorectal cancers. These data demonstrate that the ability of LCS3 to cause cell death is not unique to LUAD and may justify investigation in cancers of other histologies.

### LCS3 induces ROS and NRF2 pathway activation in sensitive LUAD cells

To understand the molecular processes governing the cellular response to LCS3, we profiled the transcriptome and proteome by RNA microarray and by stable isotope labeling of amino acids in cell culture (SILAC) (15), respectively. To assess temporal changes in RNA abundance induced by LCS3, two sensitive LUAD cell lines—H23 (KRAS mutant) and H1650 (EGFR mutant)— were treated with 3µM LCS3 for 3, 6 or 12 hours and profiled at each time point with Affymetrix RNA microarrays (Fig. 2a, Supplementary Fig. 2a). We combined genes that were significantly upregulated by LCS3 relative to the vehicle control at all time points in H23 and H1650 to determine a core set of 71 LCS3-modulated genes. Using this gene set, we queried the gene ontology (GO) knowledge base for biological processes that are affected by LCS3 and used the computational systems biology research tool Enrichr to predict the upstream regulators that control the transcriptional response to LCS3 (16, 17). Similarly, we performed a GO analysis on the set of proteins that were significantly upregulated by LCS3 at 24 hours in a SILAC study (Supplementary Fig. 2b). The GO analyses of the transcriptome and proteome expression datasets suggest that LCS3 elicits a cellular response to oxidative stress, including pathways responsive to increased levels of reactive oxygen species (ROS) (Fig. 2b). ROS include radicals, ions, or molecules that have an unpaired electron in the valence electron shell, including superoxide, hydroxyl radicals, hydrogen peroxide, peroxyl radicals, and other highly reactive species (18). It is widely recognized that LUAD cells produce increased levels of ROS relative to non-malignant cells, due to irregular mitochondrial, metabolic, and oxidase activities, as well as hyperactive growth signaling (18, 19). LUAD cells often lack the means to eliminate ROS; thus they sustain elevated levels of ROS and require the expression of cytoprotective genes to survive (18).

**Fig. 2.**
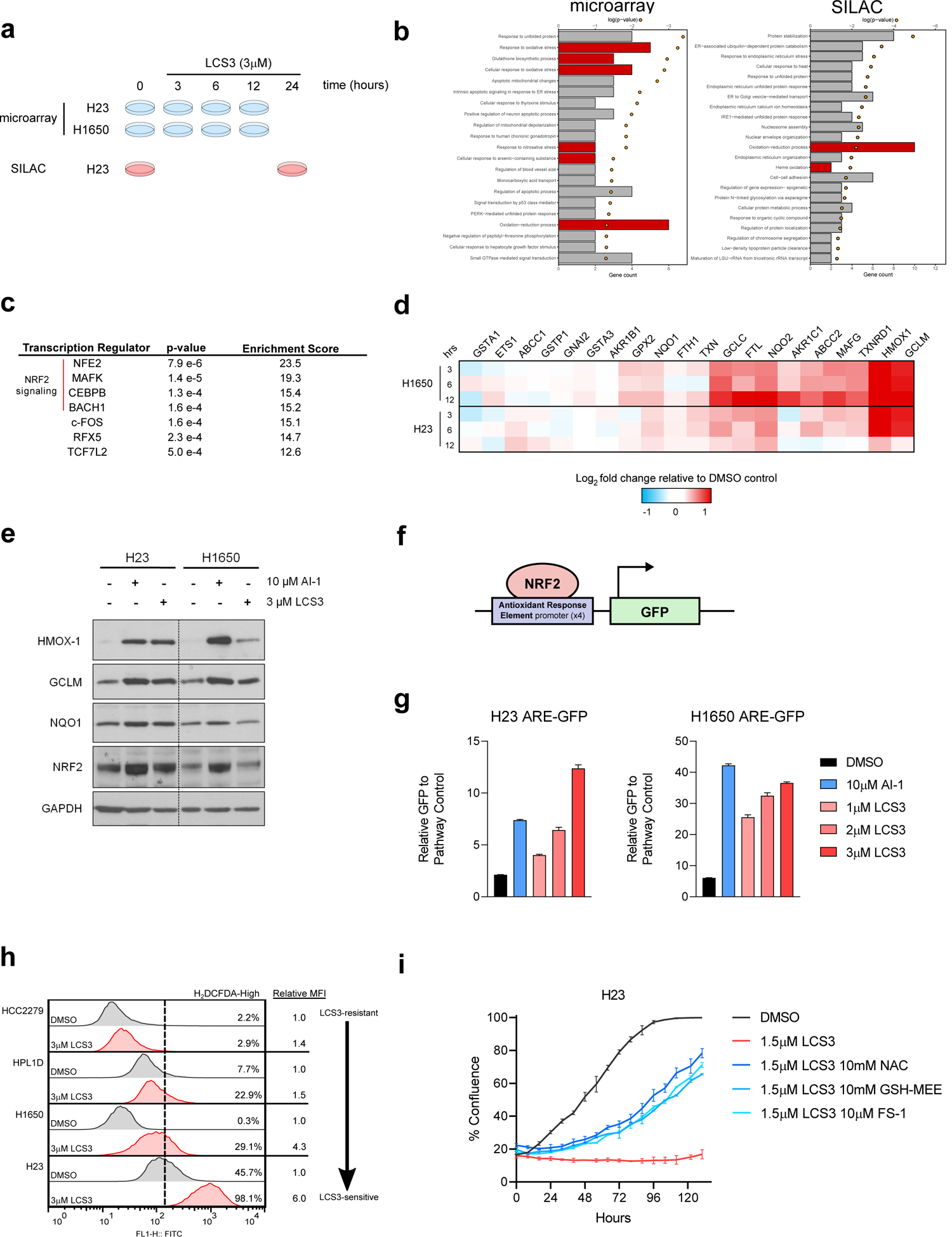
LCS3 induces ROS and NRF2 pathway activation. **a**, Experimental design schematic of LUAD cells treated with LCS3 and expression profiled by microarray and SILAC. **b,** Gene ontology analysis of biological processes upregulated by LCS3 determined by transcriptome and proteome profiling. Red bars signify a direct relation to redox homeostasis. **c,** Enrichr prediction of upstream transcription regulators of LCS3-induced RNA expression changes. **d,** Heatmap of NRF2 transcriptome signature in response to LCS3. **e,** Western blot showing induction of NRF2-regulated proteins by LCS3 and KEAP1 inactivator AI-1 (positive control). **f,** Schematic of antioxidant response element-green fluorescent protein (ARE-GFP) reporter which was transduced by lentivirus into H23 and H1650 cell lines. **g,** GFP expression of ARE-GFP relative to pathway control in response to AI-1 and LCS3. Data shown are the mean + SEM. **h,** Flow cytometry with oxidative stress sensor H_2_DCFDA detects induction of ROS after 90 minutes of treatment. **i,** Growth rescue experiment of LCS3-induced cell death in sensitive cell line H23 by antioxidants. Error bars are +/- SEM.

In agreement with the GO analysis, Enrichr analysis demonstrated that the top four predicted upstream effectors of LCS3-induced transcriptome changes are all proteins that are crucial to the cellular response to oxidative stress (Fig. 2c). These regulators, including MAFK, CEBPB, and BACH1, each function as cofactors in the transcriptional activation of nuclear factor erythroid 2-related factor (NRF2), which serves as the master regulator of the cellular response to ROS (20, 21). Correspondingly, we directly examined an established NRF2 gene expression signature (22) at each time point during LCS3 treatment in H23 and H1650 cell lines and observed an induction of NRF2-regulated genes (Fig. 2d). In a normal cellular state, NRF2 is sequestered and polyubiquitinated by the constitutively active E3 ubiquitin ligase Kelch-like ECH-associated protein 1 (KEAP1), accelerating NRF2 turnover (20). Upon oxidative or electrophilic stresses, crucial redox-sensitive KEAP1 cysteine residues become oxidized, impairing the ability of KEAP1 to repress NRF2 (21). The resulting stabilization of NRF2 facilitates its translocation to the nucleus, localization to the antioxidant response element (ARE), and transcription of cytoprotectant genes (23, 24) (Supplementary Fig. 2c).

To confirm the activation of NRF2, we treated H23 and H1650 cell lines with 3µM LCS3 and assessed the expression of NRF2 targets by western blot to measure the abundance of the protein products. We found noticeable increases in the protein levels of NRF2 and of the products of selected downstream targets of NRF2 in both cell lines (Fig. 2e). To determine if LCS3 induces enhanced binding of NRF2 to its consensus DNA sequence—the ARE (23)—upon LCS3 treatment, we introduced an ARE-green fluorescent protein (GFP) reporter into H23 and H1650 cell lines (Fig. 2f) and found that LCS3 induced the production of GFP in both cell lines in a dose-dependent manner. The response was similar to that seen with AI-1, a compound that enhances NRF2 levels by covalently inactivating KEAP1 (24) (Fig. 2g).

Since NRF2 is a principal regulator of redox homeostasis and the response to oxidative stress, we asked if LCS3 increased ROS levels in lung cells; we found that ROS accumulated in LCS3-sensitive cancer cell lines H23 and H1650, but not in the resistant HCC2279 cancer cell line or the non-transformed HPL1D lung cell line (Fig. 2h). We next sought to determine if antioxidant compounds would rescue LCS3-induced cell death and showed that three antioxidant agents—N-acetylcysteine (NAC), glutathione monoethyl ester (GSH-MEE), and ferrostatin-1 (FS-1)— reduced the toxicity of LCS3 in H23 cells (Fig. 2i). Collectively, these findings indicate that LCS3-sensitive LUAD cell lines respond to LCS3 by accumulating ROS and activating the NRF2 transcription program.

### Thermal proteomic profiling identifies redox regulating proteins as potential LCS3 targets

After establishing that the cellular response to LCS3 includes apoptosis and oxidative stress, we sought to identify molecular targets of LCS3 through a proteomics-based approach. We used TPP to identify proteins in H23 cell lysates that formed thermally-stable complexes with LCS3 (Fig. 3a) (12). A total of 5,593 proteins were detected by mass spectrometry in the lysates, and we employed the non-parametric analysis of response curves (NPARC) (25) to rank them according to their ability to form heat-stable complexes with LCS3. This strategy prioritized 77 proteins involved in diverse cellular functions, including cell structure and motility, RNA processing, and metabolism (Figs. 3b and 3c).

**Fig. 3.**
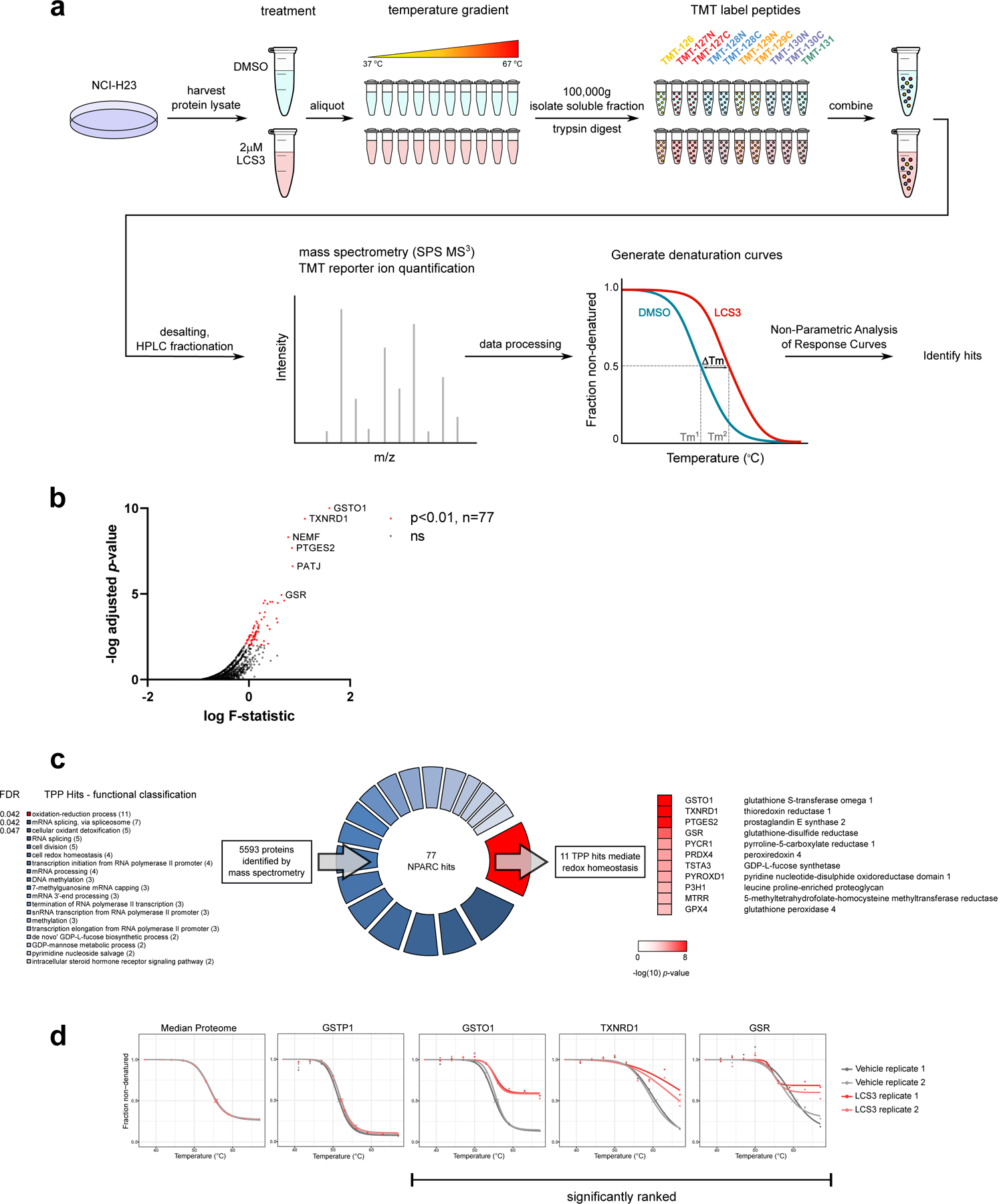
Thermal proteome profiling (TPP) identifies candidate LCS3-interacting proteins that mediate redox homeostasis. **a**, Schematic workflow of TPP methodology. Significant non-parametric analysis of response curves (NPARC) ranking indicates a protein-drug interaction. **b,** TPP NPARC identification of proteins with high F-statistic and *p*-value < 0.01 identify LCS3-interactor candidates. **c,** Functional categorization of TPP hits by DAVID Bioinformatics Database GO analysis. The false discovery rate adjusted *p*-values (FDR) of significant biological processes are shown. **d,** Select thermal denaturation curves.

The ‘oxidation-reduction process’ GO category contained the most proteins on this list and was ranked the most significant; in view of the oxidative stress observed after addition of LCS3, the effector target protein(s) seemed likely to belong to this functional class. The three most highly ranked candidates within this category were glutathione-*S*-transferase omega-1 (GSTO1; ranked 1^st^ overall), TXNRD1 (ranked 2^nd^), and GSR (ranked 6^th^); all three showed greater thermal stabilization relative to the median proteome than most other proteins (Fig. 3d). GSTO1 is known to play a protective role against oxidative stress by adding glutathione to protein thiols that are otherwise vulnerable to irreversible oxidation (26). The disulfide reductases TXNRD1 and GSR are central to the thioredoxin and glutathione antioxidant systems, which perform diverse cellular roles; most notably, they facilitate the reduction of ROS and oxidized cellular proteins (27). These and other proteins ranked with high significance by TPP; each of them function uniquely in the maintenance of redox homeostasis, may plausibly interact with LCS3, and deserve further investigation as potential drug targets.

### LCS3 inhibits the enzymatic activity of TXNRD1 and GSR

We examined the ability of LCS3 to interact with the top-ranked TPP hit, GSTO1. We were unable to obtain purified enzymatically active GSTO1 to assay its activity *in vitro* in the presence of LCS3; however, previous work has shown that GSTO1 inhibitors can be screened by their ability to bind to the catalytically active site. The cell-permeable fluorescent probe 5-chloromethylfluorescein diacetate (CMFDA) irreversibly binds the GSTO1 active site, and competition for the site by small molecules diminishes in-gel fluorescence (28). We performed an in-gel fluorescence assay by pre-incubating purified GSTO1 with LCS3 or the known GSTO1 inhibitor C1-27 as a positive control. C1-27, but not LCS3, prevented binding of CMFDA, implying that LCS3 does not occupy the GSTO1 active site and is thus unlikely to be an inhibitor of the enzyme (Supplementary Fig. 3a). We also tested the responses of LCS3-sensitive and LCS3-resistant cell lines to C1-27 and did not observe any differential sensitivity to C1-27 among cell lines (Supplementary Fig. 3b), suggesting that LCS3 and C1-27 exert toxicity by different means.

The results with GSTO1 prompted the examination of the other TPP screen candidates as potential targets of LCS3. The second-ranked candidate from the TPP study, thioredoxin reductase 1 (TXNRD1), is structurally and functionally similar to another candidate, glutathione disulfide reductase (GSR—ranked 6^th^). Together, these enzymes are essential components of the two systems that regulate the cellular redox state and other diverse biological processes through thioredoxin and glutathione. TXNRD1 and GSR possess NADPH-dependent disulfide reductase activity and reduce thioredoxin (TRX) and glutathione disulfide (GSSG) substrates, respectively. After reduction, TRX and GSH participate in many reactions that scavenge ROS and reduce cellular substrates (29). The antioxidant activity of TRX is primarily through reducing peroxiredoxin proteins, which are among the most abundant proteins in mammalian cells (30). In addition, reduced glutathione (GSH) is a versatile adaptor molecule and an essential cofactor for enzymes that require thiol-reducing equivalents, particularly in those with redox-active cysteines in their catalytic sites (31). Further, we showed that the monoethyl ester form of glutathione, GSH-MEE, with its efficient uptake by cells, was able to partially rescue cells from LCS3-induced toxicity, suggesting that GSH levels may be important in regulating the response to LCS3 (Fig. 2i).

We next determined whether GSR and TXNRD1 are direct binding targets of LCS3 by performing *in vitro* assays with purified protein. Both enzyme reactions catalyze the transfer of electrons from NADPH to either GSSG or Ellman’s reagent (DTNB), which has a disulfide bond that can be reduced by TXNRD1, replacing the requirement for a purified TRX substrate (32, 33). Both GSR and TXNRD1 were inhibited by LCS3 (IC_50_ = 3.3µM and 3.8µM, respectively) (Fig. 4a). We also sought to establish the mechanism of enzymatic inhibition by LCS3. Pre-incubation of an enzyme with a covalent inhibitor increases enzymatic inhibition (indicated by a lower IC_50_) and we used this strategy to assess the reversibility of the LCS3-protein interaction. As a positive control, we pre-incubated GSR with the covalent disulfide reductase inhibitor 2-AAPA (50µM); as anticipated, the longer the pre-incubation period, the greater the enzymatic inhibition. In contrast, pre-incubation with 3µM LCS3 did not affect the amount of inhibition, suggesting that LCS3 inhibits GSR by a reversible mechanism (Fig. 4b).

**Fig. 4.**
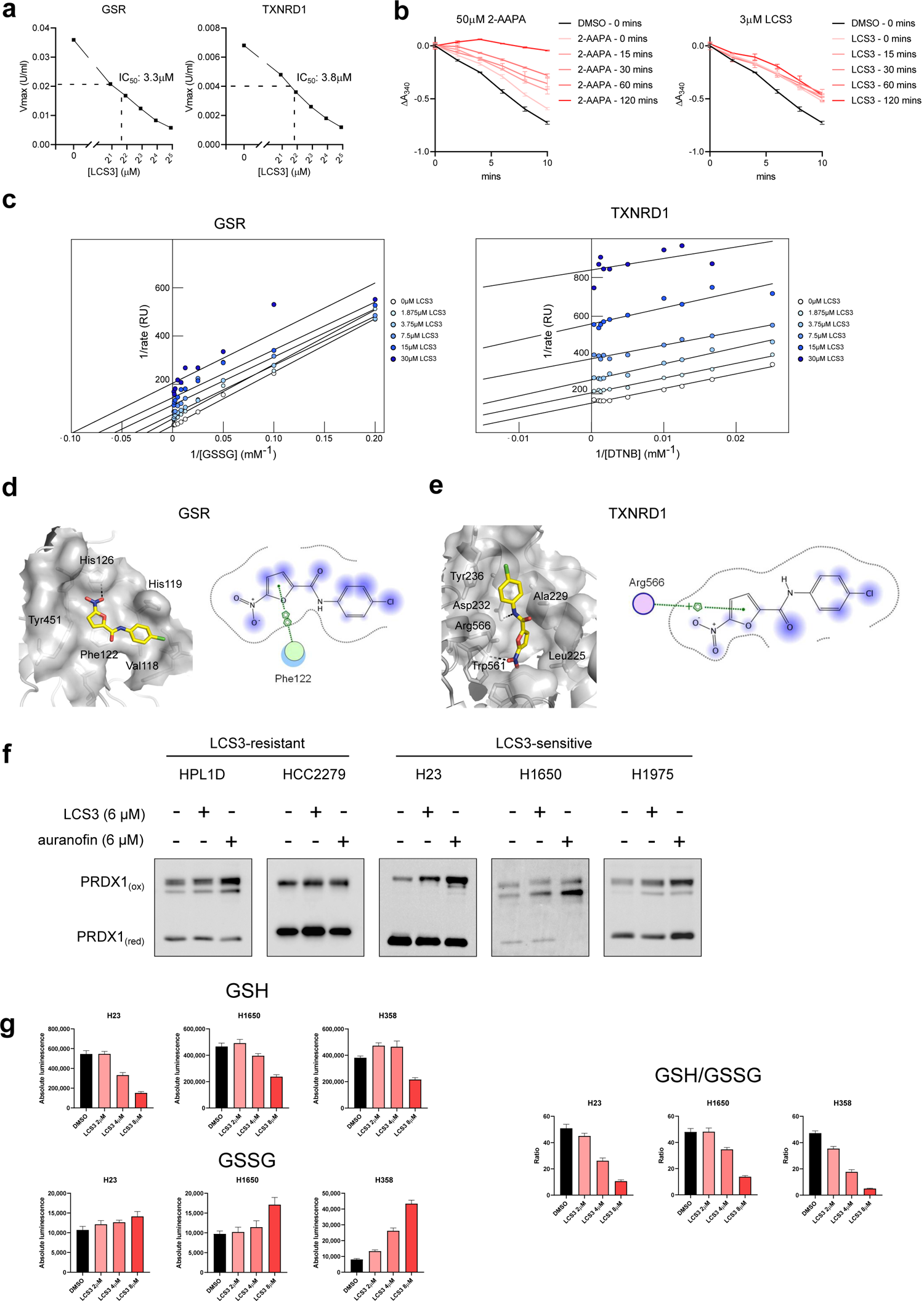
Validation of GSR and TXNRD1 as putative LCS3 targets. **a**, *In vitro* enzymatic assays of purified GSR and TXNRD1 in the presence of LCS3. **b,** Pre-incubation assay with purified GSR and specified pre-incubation durations with covalent GSR inhibitor 2-AAPA (positive control) and LCS3. Error bars are +/- SEM. **c,** Lineweaver-Burk plots of GSR and TXNRD1 inhibition by LCS3. **d-e,** *In silico* molecular docking of LCS3 into the homodimer interfaces of GSR and TXNRD1. Schematic of LCS3 arene-arene and arene-cation interactions with amino acid residues of GSR and TXNRD1, respectively. **f,** PRDX1 redox status upon LCS3 and auranofin treatment determined by redox western blot. **g,** Metabolite assay to determine the relative concentrations of GSH and GSSG in LUAD cell lines. Error bars are +/- SD.

To understand how LCS3 affects the enzyme kinetics of GSR and TXNRD1, we generated Lineweaver-Burk plots (Fig. 4c). Plotting the reciprocal of enzymatic rates by the reciprocal of substrate concentrations for each concentration of LCS3 produced lines of best fit that are parallel—a strong indication that LCS3 acts on both GSR and TXNRD1through an uncompetitive inhibitory mechanism. This suggests that LCS3 does not compete directly with the cellular substrates of these enzymes in their free state, but rather inhibits the enzymes following the formation of an enzyme-substrate complex (34). These data suggest that LCS3—as an uncompetitive inhibitor—preferentially binds the disulfide reductases at high concentrations of oxidized substrates, GSSG or DTNB, and does not bind the enzymes in the absence of substrate.

Based on these observations, we next performed *in silico* molecular docking studies to predict how LCS3 physically interacts with GSR and TXNRD1. The molecular docking simulation predicts that the nitro group of LCS3 forms a crucial hydrogen bond with GSR at His126 (Fig. 4d). Moreover, the furan moiety of LCS3 is predicted to form an arene-arene electrostatic interaction with nearby Phe122. Additional hydrophobic contacts were predicted between LCS3 and Val118, His119 and Tyr451. Enzymatic activity of GSR is dependent on its homodimer formation, in which a side-chain extending from each subunit binds and processes the GSSG substrate. This homodimerization is indispensible for GSR activity and a strong hydrogen bond interaction has been described between the imidazole rings of His119—His126’ (35). The molecular docking data suggest that the LCS3 nitro subgroup displaces the His119— His126’ hydrogen bond between subunits. We theorize that this displacement, in addition to the other LCS3-GSR interactions noted earlier, confer the ability of LCS3 to inhibit enzymatic activity.

In a similar fashion, TXNRD1 homodimerizes to perform its catalytic function and LCS3 is predicted to strongly bind the side chain residues responsible for homodimer formation. The nitro group of LCS3 is predicted to form a hydrogen bond with Trp561 of TXNRD1 and LCS3 is also predicted to interact via hydrogen bond formation with Asp 232 and an arene-cation interaction with Arg566 (Fig. 4e). Critical hydrophobic contacts are predicted between LCS3 and Leu225, Ala229 and Tyr236. Arg566 is situated at the homodimer interface on a segment which functions as a guiding bar to stabilize the flexible C-terminal arm during electron transfer to the TRX substrate (36). Arg566 has been reported to form hydrogen bonds with Asp232 (37), and the arene-cation and hydrogen bond formation of LCS3 with both these residues may disrupt this structural interaction. Together, the molecular docking data predict that LCS3 forms molecular bonds with residues in the homodimer interfaces of both GSR and TXNRD1 and suggest that the reversible occupancy of these intermolecular cavities by LCS3 disrupts the molecular functions of these disulfide reductases.

In light of the docking analysis that indicated how LCS3 likely interacts with GSR and TXNRD1, we probed the structure-activity relationship (SAR) between the chemical structure of LCS3 and its cytotoxic activity. We conducted a cell-based sensitivity screen to investigate how LCS3 and 26 structural analogues inhibit the growth of a panel of lung cell lines (Table S1). We found that only compounds that possess the nitro functional group exhibit toxicity in LUAD cells (Supplementary Fig. 3c), demonstrating the necessity of this moiety for the inhibition of cell growth caused by LCS3. We next evaluated if the nitro subgroup is essential for enzymatic inhibition of GSR by an *in vitro* enzymatic assay with purified protein. We found that a high concentration of LCS3 (20µM) abolished all GSR activity, but the same concentration of an LCS3 analogue (N-(4-chlorophenyl)furan-2-carboxamide; PubChem ID CID790315) lacking the nitro subgroup did not inhibit GSR (Supplementary Fig. 3d). Thus, these data suggest that the nitro functional group of LCS3 is essential for its ability to inhibit enzymatic activity and subsequently elicit death of cancer cells. These data also support the molecular docking analysis, which predicts that the nitro subgroup of LCS3 forms crucial hydrogen bonds with GSR His126 and TXNRD1 Trp561.

We then assessed whether LCS3 inhibits the downstream products of GSR and TXNRD1 activity within cells. We performed a redox western blot, which preserves protein oxidation, to determine the relative oxidation state of peroxiredoxin 1 (PRDX1), a direct substrate of TRX and a surrogate for the detection of TRX activity (38, 39). Auranofin, a covalent inhibitor of TXNRD1, served as a positive control. Addition of LCS3 markedly increased oxidation of PRDX1 in LCS3-sensitive LUAD cell lines after 2 hours, to a degree comparable to that produced by the known TXNRD inhibitor, auranofin; but LCS3 did not increase oxidation of PRDX1 in the insensitive cell lines HPL1D or HCC2279 (Fig. 4f).

To assess the effects of LCS3 on GSR, we determined the intracellular ratio of GSH to GSSG after exposing LUAD cell lines to 2-8µM LCS3. Addition of LCS3 decreased the concentration of GSH and increased the concentration of GSSG in LCS3 sensitive cell lines, resulting in a dose-dependent decrease in the [GSH]/[GSSG] ratio (Fig. 4g). These data suggest that LCS3 interacts with GSR and TXNRD1, inhibits their enzymatic activities, and reduces the levels of their products in LCS3-sensitive LUAD cells.

### LCS3 induces synergistic inhibition of the glutathione and thioredoxin pathways

After demonstrating that LCS3 inhibits the disulfide reductases GSR and TXNRD1, we aimed to establish whether loss of these enzymatic activities is toxic to LUAD but not non-cancerous lung epithelial cells. It is well-established that combined inhibition of the glutathione and thioredoxin pathways synergizes to induce cell death in some types of cancer cells (40–46). We hypothesized that if LCS3 exerts cytotoxicity by synergistic inhibition of these two compensatory pathways, then combining LCS3 with an agent that inhibits either pathway should further sensitize LUAD cells to LCS3. This assumes that co-treatment of LCS3 with a compound that inhibits one of its effectors will reduce the concentration of LCS3 required to inhibit both pathways and elicit a lethal phenotype.

Using all three dual pairings, we tested the combinatorial effects of LCS3, the TXNRD1 inhibitor auranofin, and the GSH synthesis inhibitor buthionine-sulfoximine (BSO) on the growth of LUAD cell lines (Fig. 5a and Supplementary Fig. 4a). Consistent with other reports, we found the combination of auranofin and BSO to be highly synergistic in LUAD cell lines, demonstrating the compensatory nature of the glutathione and thioredoxin pathways. The effects of LCS3 were also synergistic with either BSO or auranofin (Fig. 5a), supporting the case that LCS3 exerts toxicity by disrupting both redox pathways. Additionally, we found that siRNA-mediated knockdown of GSR (siGSR) sensitized cells to LCS3, which was also observed when combining auranofin and siGSR—this shared property of sensitizing LUAD cells to GSR depletion further suggests at least a partial overlap exists between the mechanisms of LCS3 and TXNRD1 inhibitor auranofin (Supplementary Fig. 4b). However, we did not observe additive effects when using BSO and siRNA targeting of TXNRD1 (Supplementary Fig. 4c), suggesting that other thioredoxin reductases may compensate for loss of TXNRD1, but not for the inhibitory effects of LCS3 or auranofin. These data are supportive of a mechanism by which LCS3 disables both the glutathione and thioredoxin pathways; in this way, increased reliance on either of the two pathways, after addition of pathway specific inhibitors or siRNAs, impairs the ability of LUAD cells to maintain redox homeostasis and consequently sensitizes cells to LCS3.

**Fig. 5.**
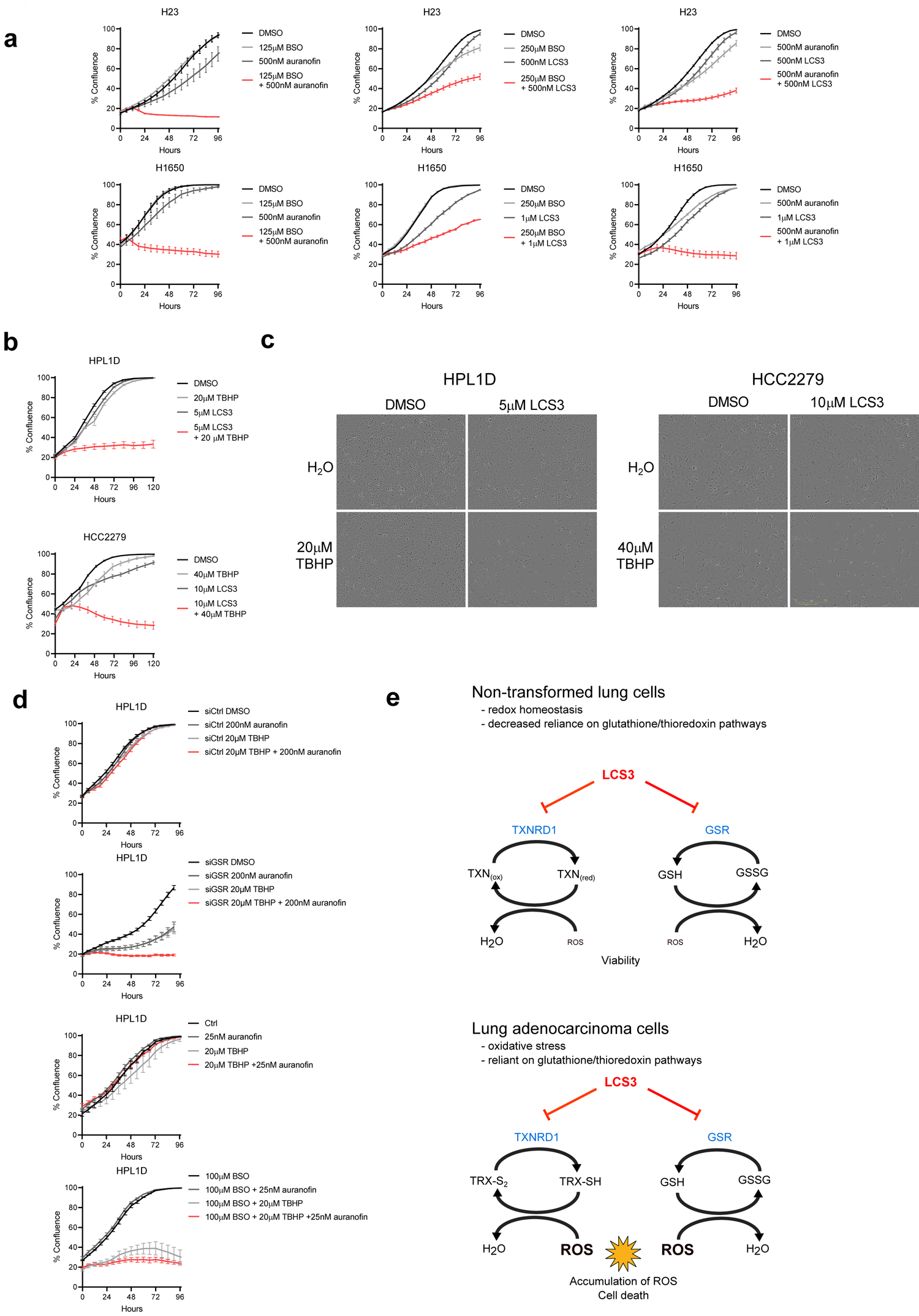
Sensitization of LUAD cells to ROS by dual inhibition of glutathione and thioredoxin pathways. Incucyte growth assay results are shown as mean +/- SEM. **a,** LCS3 and inhibitors of the glutathione and thioredoxin pathways (BSO and auranofin, respectively) in combination. **b,** Treatment with LCS3 alone and in combination with TBHP in LCS3-resistant cell lines. **c,** Incucyte images taken at 72 hours of HPL1D and HCC2279 cells treated with LCS3 alone or in combination with TBHP. **d,** TBHP affects viability in HPL1D upon dual inhibition of glutathione and thioredoxin pathways achieved by auranofin and either GSR knockdown or BSO. **e,** A working model for how LCS3-mediated dual inhibition of glutathione and thioredoxin pathways elicits accumulation of ROS and cell death in LUAD cells.

We have shown that LCS3-sensitive LUAD cell lines are sensitive to combined inhibition of glutathione and thioredoxin pathways, and noted that insensitive cell lines have low basal levels of ROS (Fig. 2h). Consequently, we asked whether supplying ROS to insensitive cell lines would increase reliance on glutathione and thioredoxin pathways and thereby sensitize resistant cells to LCS3. We treated LCS3-resistant cell lines HPL1D and HCC2279 with LCS3 alone and in combination with tert-butyl hydroperoxide (TBHP), a direct-acting hydroperoxide source of exogenous ROS. These cell lines were insensitive to LCS3 or TBHP alone, but growth was strongly inhibited when the agents were combined (Fig. 5b and Fig. 5c), implying that LCS3-resistant cells can be sensitized to LCS3 by the addition of ROS. It appears that, when challenged with oxidative stress, these LCS3-resistant cell lines become dependent on the glutathione and thioredoxin pathways to maintain control of redox balance; dual inhibition of these pathways by LCS3 allows ROS to accumulate and compromises cell survival.

We next evaluated if sensitization of LCS3-resistant cells to ROS could be phenocopied with known inhibitors of the glutathione and thioredoxin pathways. When HPL1D cells were grown with TBHP and a combination of auranofin and either BSO or siGSR, their growth was inhibited, although they were tolerant of auranofin, TBHP, or GSR depletion alone. Combination of TBHP and BSO induced a degree of toxicity; however, this effect was increased with the addition of auranofin (Fig. 5d). These data are consistent with our prior observation that TBHP sensitizes LCS3-resistant cells to LCS3 and suggest that inhibition of both GSR and TXNRD1 contributes to LCS3-induced toxicity. Therefore, cells exhibiting higher basal levels of ROS— including the majority of LUADs—are more likely to be sensitive to LCS3-mediated inhibition of the glutathione and thioredoxin pathways, as shown schematically in Fig. 5e.

### CRISPR-Cas9 knockout screen confirms the role of LCS3 in mediating antioxidant response and reveals suppression of NQO1 as a mechanism of resistance

To complement the effort to identify targets of LCS3 with TPP and to interrogate potential mediators of the LCS3 response systematically, we performed a genome-wide, loss-of-function CRISPR-Cas9 screen in LCS3-sensitive LUAD cells. H358 cells were infected with a lentiviral pool generated using the Toronto KnockOut Library v3 (TKOv3) library, which contains Cas9 in conjunction with 70,948 gRNAs targeting 18,053 protein-coding genes (47). Transduced cells were passaged in the presence of DMSO, 4µM LCS3, or 8µM LCS3 for 14 population doublings. DNA from the three cultures was assessed for the degree of enrichment or depletion of the sgRNAs in the original library in response to either concentration of LCS3 (Fig. 6a). The sgRNAs targeting the genes encoding NQO1 and glutathione-*S*-transferase P (GSTP1) were significantly enriched, while those targeting glutamate-cysteine ligase (GCLM) and KEAP1 were significantly depleted, in cultures grown at both concentrations of LCS3.

**Fig. 6.**
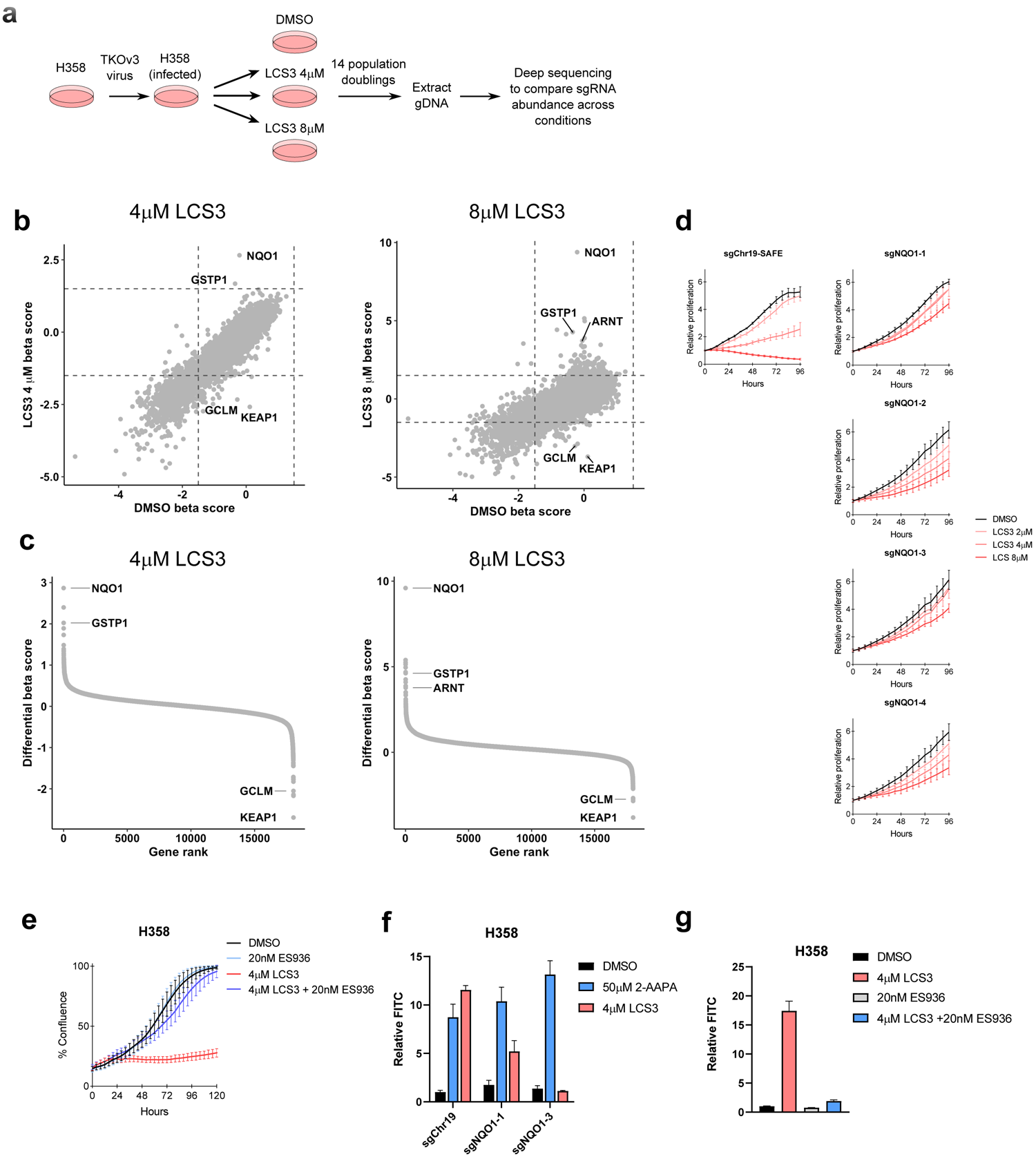
Genome-wide CRISPR-Cas9 knockout screen identifies NQO1 suppression as a mechanism of resistance. Incucyte growth assay results are shown as mean +/- SEM. H_2_DCFDA assay results are shown as mean +/- SD. **a,** Schematic of Genome-wide loss-of-function CRISPR-Cas9 knockout screen. **b,** Scatterplots of CRISPR screen beta scores of LCS3 treatments and vehicle control. **c,** Rank of proteins as determined by treatment beta scores subtracted by control treatment scores. **d,** Incucyte growth experiment of H358 sgNQO1 cells upon LCS3 treatment compared to control. **e,** Growth rescue experiment of LCS3-induced cell death in sensitive H358 by NQO1 inhibitor ES936. **f,** Flow cytometry with oxidative stress sensor H_2_DCFDA upon LCS3 and 2-AAPA (positive control) in H358 sgNQO1 cell lines. **g,** Flow cytometry with oxidative stress sensor H_2_DCFDA upon combination treatment with LCS3 and NQO1 inhibitor ES936.

It is notable that many proteins encoded by the affected genes play prominent roles in regulating redox balance (Fig. 6b) (20, 48). The sgRNAs targeting the genes encoding NQO1 and KEAP1 were, respectively, the most significantly enriched and the most significantly depleted under both LCS3 conditions. NQO1, an NRF2 target gene, is a NADPH dehydrogenase that reduces quinones during detoxification and is involved in the cellular adaptation to stress (49); however, it has been reported that NQO1 activity can elicit a bioreductive activation of substrates that is associated with the production of ROS and exacerbation of redox imbalance (50). While loss of NQO1 had no effect on cellular fitness in H358 cells under normal growth conditions, it greatly increased the relative ability of cells to proliferate in the presence of LCS3 (Fig. 6b). We confirmed this observation by establishing stable cell lines in which the NQO1 gene was knocked out or by inhibiting NQO1 with the small molecule ES936; both approaches demonstrated nearly complete protection against LCS3-mediated toxicity (Fig. 6d and 6e). In addition, we observed a dramatic decrease in the induction of ROS by LCS3 in H358 cells in which NQO1 was suppressed by either sgNQO1 or ES936, mirroring the reduction in toxicity (Fig. 6e and 6f). In addition, cells in which the NQO1 gene was inactivated by sgNQO1 did not activate the NRF2 pathway relative to the control cell line (Supplementary Fig. 5a). Thus, the phenotype of NQO1-null cells—loss of LCS3-induced, ROS-mediated cell toxicity—is not likely mediated by changes in regulation of NRF2. In H358 sgChr19-SAFE control cells, but not H358 sgNQO1 cells, LCS3 induces an increase in NADPH, further suggesting that the cellular effects of LCS3 are inhibited upon knockout of NQO1 (Supplemental Fig. 5b).

The depletion of *KEAP1* sgRNAs, implying selection against *KEAP1*-null cells during growth in LCS3, was unexpected. KEAP1 is a negative regulator of NRF2, and elevated NRF2 activity would be predicted to allow cells to better tolerate the induction of ROS by LCS3. To test this, we used the CRISPR-Cas9 method to inactivate *KEAP1* in H358 cells with multiple sgRNAs and found no significant effects on sensitivity of cell growth to LCS3, despite an increase in NRF2 levels (Supplemental Fig. 5c and 5d). This finding appears to conflict with the results of the knockout screen. However, NRF2 controls the transcription of *NQO1* and *GSTP1* (51), genes that were depleted in cells that grew preferentially in the presence of LCS3.

Therefore, it is possible that the selection against *KEAP1* sgRNAs in the context of the screen is related to the induction of these and other NRF2 targets. Indeed, cells bearing sgRNAs for the transcription factor *ARNT*—which, together with NRF2, controls *NQO1* expression by binding to the Xenobiotic Response Element motifs in response to electrophilic stress (52)—were also positively selected in 8µM of LCS3, which further supports this theory (Fig. 6b). The results of the CRISPR screen support a framework in which LCS3 disrupts redox homeostasis in sensitive LUAD cell lines and highlights NQO1 activity as a key determinant of LCS3 sensitivity and a potential biomarker for response.

This view is further supported by additional results from the CRISPR screen. GCLM catalyzes the ligation of cysteine to glutamate in the first and rate-limiting step of glutathione biosynthesis (40, 53); inactivation of *GCLM* is predicted to impair the downstream glutathione pathway and yield an effect similar to the knockdown of GSR or to the addition of BSO—factors that cooperate with LCS3. Therefore, the selection against the suppression of GCLM in the CRISPR screen is consistent with the observed synergies between LCS3 and other glutathione pathway disruptors.

The sgRNAs for glutathione synthetase (GSS), which regulates the second step of glutathione biosynthesis (53, 54), were also significantly depleted in the CRISPR screen, but only in the cultures grown in 8µM of LCS3. This finding further supports a mechanism by which LCS3 elicits cancer cell death via the disruption of the glutathione and thioredoxin pathways. Another potentially informative gene depleted in the CRISPR screen, *GSTP1*, encodes a xenobiotic detoxification enzyme which conjugates reduced glutathione to electrophilic compounds to facilitate excretion of GSH (55). Thus, sgRNA-mediated inactivation of *GSTP1* would plausibly promote cell survival in the context of LCS3 treatment by increasing the availability of free GSH.

## Discussion

We investigated the mechanism of action and effector targets of the novel small molecule LCS3, which was uncovered previously in a high-throughput, phenotype-based drug screen designed to identify compounds that selectively inhibit LUAD growth (10). Here we have examined the activity of LCS3 in an expanded panel of LUAD and normal lung cells and employed a chemical biology approach to seek new druggable targets and cell survival dependencies that could potentially be exploited for the treatment of lung cancer. Using an integrative gene expression and proteomic analysis of treated cells, we found that LCS3 inhibits LUAD through an oxidative stress-induced mechanism of apoptotic cell death. TPP and enzymatic assays revealed that the disulfide reductases GSR and TXNRD1 are targets of LCS3 and we showed that inhibition of both proteins synergistically induces cell death.

Failure to identify or characterize the molecular targets of novel anti-cancer compounds has been a major limitation of phenotype-based chemical library screens. One of the most widely used approaches in target discovery is affinity proteomics, which involves the covalent linking of a derivatized small molecule to an immobilized structure—such as magnetic or carbohydrate beads—coupled with affinity chromatography and mass spectrometry (11). The principal disadvantage with this approach is that it involves modifying the native structure of the compound of interest and the resulting shift in chemical properties may impact drug-protein interactions (56). As opposed to traditional affinity-based methods, TPP offers the potential to efficiently identify proteins that bind compounds that are therapeutic candidates, without the need for chemical derivatization; in this way, TPP limits false-positive findings and streamlines the workflow (57). While another group recently determined a target of the autophagy inhibitor indophagolin using a TPP-based strategy (58), to our knowledge few other studies have used TPP to identify the effector targets of any small molecule inhibitor—rather than off-target binders of well-characterized compounds—in human cells. Therefore, our multifaceted approach—combining systematic interrogation of the cellular response to drug treatment with a comprehensive assessment of protein targets in drug-sensitive cells with TPP—may serve as a template for characterizing the mechanism of action of novel anti-cancer compounds in the future.

Increased production of ROS is associated with oncogene signaling and transformation in lung and other solid cancers (40); hence targeting redox homeostasis has become an attractive approach for the development of new cancer therapies. LUAD cells commonly have genetic alterations that lead to NRF2 pathway activation and increased cellular cysteine through upregulation of system Xc-in order to synthesize GSH and other ROS-scavenging metabolites (59). Demonstrating the importance of these programs for cell fitness and survival, strategies that inhibit key NRF2 effectors or cause cysteine depletion can induce apoptosis and other forms of cancer cell death both *in vitro* and *in vivo* (60, 61). Indeed, another compound, known as LCS1, from the original screen in which LCS3 was identified, was subsequently found to inhibit SOD1, a key NRF2 target gene product that mitigates ROS by transforming superoxide anions into hydrogen peroxide (11). Our findings in this report build on the concept that at least some cancer cells are exquisitely sensitive to oxidative stress and demonstrate a reliance on the glutathione and thioredoxin systems to maintain redox homeostasis and survival. While normal cells can tolerate LCS3 because of low basal levels of ROS, adding exogenous ROS to mimic the oncogenic state leads to LCS3 susceptibility and lethality (Fig. 5b), suggesting that cells in a state of oxidative stress have increased dependence on the glutathione and thioredoxin systems. Furthermore, in LUAD cells, disruption of either of these pathways alone does not have a major effect on cell viability; however, their combined inhibition in the presence of ROS leads to synthetic lethality. This context-dependent sensitivity to LCS3 indicates that strategies that target the glutathione and thioredoxin pathways may yield a favorable therapeutic ratio, with selective cytotoxicity in cancer cells.

The concept of dual targeting of the glutathione and thioredoxin pathways in cancer has gained attention in the last decade (40–46). While the complete mechanism of action resulting from the combined inhibition of these pathways remains unclear, a discernible synergy arises from their interconnection and compensatory nature. As an example, glutaredoxin-2 (GLRX2) is reduced non-enzymatically by GSH but can be reduced by thioredoxin reductases when GSH is depleted (62). In a reciprocal fashion, glutaredoxin-1 (GLRX1) can rescue TXNRD1 loss by catalyzing the transfer of electrons from GSH to TRX (63, 64). Thus, the compensation provided by these connected pathways in alleviating oxidative stress in cancer cells is supportive of a therapeutic strategy based on dual pathway inhibition. LCS3 serves as an important proof of principle in this regard; further work based on this compound should be carried out in animal models to investigate if targeting this dependency in LUAD or in other malignancies is a viable therapeutic avenue for clinical development.

While it is known that ROS are generated as a by-product of cancer cell growth (65), a more comprehensive understanding of the redox landscape could potentially help guide rational patient selection for drugs that exploit this sensitivity (18, 19). In our lung cell line panel, HCC2279 was the sole LCS3-resistant LUAD cell line (Fig. 1b). We observed that this cell line has low basal ROS, which may explain why it is non-responsive (Fig. 2h); however, the underlying dynamics that regulate this state are unclear. Our CRISPR-Cas9 knockout screen in H358 cells, which are sensitive to LCS3 treatment, demonstrated key factors that may be useful as biomarkers for predicting sensitivity to dual glutathione and thioredoxin suppression.

Numerous sgRNAs that were enriched and depleted in cells grown in this screen disrupt genes that are involved in regulating key processes in glutathione metabolism and oxidative stress response, further supporting the mechanism of action of LCS3. However, our finding that cells containing *KEAP1* sgRNAs were depleted in the screen was unanticipated (Fig. 6c), since we predicted that activation of the NRF2 pathway would protect cells against LCS3-mediated oxidative stress. While the pleiotropic effects of NRF2 —which include activating many genes that are associated with increased sensitivity to LCS3—may explain this finding, we were unable to validate the effects of *KEAP1* disruption on LCS3 sensitivity by creating stable *KEAP1*-defective clones of H358 cells. This suggests that the temporal relationships of *KEAP1* disruption and LCS3 treatment may also be a factor, with simultaneous suppression of *KEAP1* and treatment with LCS3 having detrimental effects on cell viability. In contrast, inactivation of *KEAP1* before addition of LCS3 may permit adaptation of the relevant pathways to protect against the toxic consequences of ROS. In other words, the role of the KEAP1/NRF2 pathway in mediating sensitivity and resistance to LCS3 and other oxidative stress-inducing compounds may depend on the timing and balance of these regulatory factors.

The most striking finding from the CRISPR knockout screen was the apparent dependence of LCS3 sensitivity on NQO1 activity (Fig. 6c), supporting a key role for this enzyme in LCS3-induced cell death. We validated this finding by creating stable *NQO1* knockout cells and by using an inhibitor of NQO1; both strategies protected H358 cells from LCS3 (Fig. 6d and 6e). It is noteworthy that NQO1 has been implicated in the bioreduction of various nitroheterocyclic compounds, which is essential for inducing their anticancer activities. This process catalyzes the reduction of a nitro group of a compound into a nitro anion radical, which can subsequently react with oxygen to form a superoxide anion and hydrogen peroxide, inducing oxidative stress and toxicity in cancer cells (66, 67). As an example of this, the structurally related antibiotic nitrofurantoin has been reported to generate a nitro anion radical by redox cycling, a process which results in both the production of ROS and the regeneration of the parent molecule (68). Our SAR analysis demonstrated that the nitro moiety is essential for the anti-proliferative and enzymatic inhibition activities of LCS3, suggesting that the electron-accepting properties of the nitro group confer intermolecular affinity for the reductases GSR and TXNRD1, which are electron donors to GSSG and thioredoxin substrates. Interestingly, our *in vitro* assays with purified proteins demonstrate that LCS3 causes direct enzyme inhibition, suggesting that LCS3 activity does not require redox cycling or another form of NQO1-dependent biotransformation to inhibit GSR and TXNRD1. However, it is possible that structural analogues of LCS3 that lack a nitro group are not metabolized by NQO1 and consequently fail to generate ROS via redox cycling and thus do not exhibit activity in cell-based experiments. Thus, redox cycling mediated by NQO1, which is characteristically high in cancer cells, could explain why LCS3 induces ROS only in cancer and not in normal cells. While our demonstration that dual inhibition of the glutathione and thioredoxin pathways by BSO and auranofin recapitulates the cytotoxicity of LCS3, it is possible that NQO1 potentiates these effects through redox cycling and concomitant generation of ROS. We speculate that LCS3 may be unique in its ability both to cause oxidative stress in NQO1-expressing cancer cells and to abrogate the ability of cancer cells to cope with accumulated ROS, through parallel disruption of the glutathione and thioredoxin systems. Therefore, it is both interesting and unexpected that the highly reactive nitro group— which could be viewed as a suboptimal structural feature from a medicinal chemistry perspective—may be responsible for the selective and potent activities of LCS3. If so, further medicinal chemistry may optimize its inhibitory and pharmacological properties. Together, our data suggest that NQO1 may serve as a potential biomarker to predict LCS3 response in future preclinical studies. It might also be useful to determine which derivatives of LCS3 are produced by biotransformation and to assess their bioactivities.

Overall, we have demonstrated LCS3 is a reversible, uncompetitive inhibitor of GSR and TXNRD1 that selectively kills LUAD cells in an oxidative stress-dependent manner. Therefore, the potential utility of LCS3 lies in targeting redox homeostasis as a dependency for therapeutic purposes. Further investigations are warranted to fully explore how LSC3 and the redox pathways can be exploited for the treatment of LUAD and other cancers, especially for those with poor outcomes and limited therapeutic options.

## Methods

### Cell culture and growth conditions

All cell lines were cultured in either RPMI (Gibco, 11875119) or DMEM media (Gibco, 12430063) supplemented with 10% fetal bovine serum (Gibco, 12483020) and 1% penicillin-streptomycin (Gibco, 15140-122) and passaged by washing with Dulbecco’s phosphate buffered saline (PBS) (Gibco, 14190250), and incubating with trypsin-EDTA (0.25%) (Gibco, 25200114). Cells were maintained in a humidified incubator at 37°C and 5% CO_2_. Compounds used in this study: LCS3 (Chembridge, 5181525), NAC (Sigma, A7250-25G), GSH-MEE (Sigma, 353905), FS-1 (Sigma, SML0583-5MG), AI-1 (Sigma, SML0009-5MG), TBHP (Sigma, 458139-100ML), auranofin (Sigma, A6733-10MG), BSO (Sigma, B2515-500MG), RSL3 (Selleckchem, S8155), ES936 (Santa Cruz, 192820-78-3), 2-AAPA (Sigma, A4111-5MG), C1-27 (MedChemExpress, HY-111530).

### Cell line drug screens

High-throughput screening of LCS3 and related derivatives for SAR assessment and determination of IC_50_ values was performed as previously described (10). Cell lines were fingerprinted by STR profiling to confirm identity and mycoplasma tested prior to use. LCS3 was submitted to the Developmental Therapeutics Program at the National Cancer Institute for assessment of the NCI-60 cell line panel using an established workflow previously described (https://dtp.cancer.gov/discovery_development/nci-60/default.htm).

### Crystal violet stain

Ten thousand cells were seeded in triplicate into 6-well plates and then treated the following day with 0.1% DMSO (Fischer, BP231-100) or 1µM LCS3. After nine days, cells were washed twice in ice-cold PBS before fixed in ice-cold methanol (Fischer, A433P-4) at room temperature for 10 minutes. Cells were then stained with 0.5% (vol/vol) crystal violet solution (Sigma, HT90132-1L) at room temperature for 10 minutes, on a rotating shaker. Once stain was completed, cells were rinsed with PBS thrice and left overnight to air dry.

### Incucyte cell growth experiments

Cells were seeded in clear 96-well plates (Sigma, CLS3595) at 4,000 cells per well. One day after seeding, media containing drug treatments was added to each well and plates were placed in an Incucyte S3 live-cell imaging system within an incubator maintained at 37°C with 5% CO_2_. Images were taken at specified time intervals. For STACK experiments—H23^N^, H1650^N^, H1975^N^, H358^N^, HPL1D^N^, HCC2279^N^ cells with H2B-Cherry stably transduced by lentivirus (Gift from Torsten Wittman, Addgene #89766) (69) were seeded into 96 well plates. Twenty-four hours after seeding, cells were treated with 20nM SYTOX Green (Thermo, S7020) and 10µM RSL3 or 500nM-16µM LCS3. STACK analysis was performed by determining the lethal fraction of each cell line under each condition, determined by counting the dead cells (green and red overlap) and dividing by the total cells (red objects). Onset of cell death (D_o_) and the rate of cell death (D_R_) for each cell line were determined by fitting a plateau followed by one phase association model with GraphPad Prism 8.3.1.

Constraints used were X_o_ >1, Y_o_ >1, 0 1 and K >1. X_o_ was represented as D_o_ and K*100 was represented by D_R_. For combination experiments in Supplementary Fig. 4a, the fraction of growth inhibition was calculated by subtracting the fraction of treatment condition confluence over maximal confluence from 1 at the confluence end point, which was determined to be the first time point for each plate in which the mean control condition surpassed 97% confluence.

### Western blotting

Cells were harvested and washed with ice-cold PBS before being scraped in PBS, transferred into 15mL conical tubes and centrifuged at 1500rpm for 5min. Cells were resuspended in RIPA lysis buffer (Pierce Protein Biology, P123227) supplemented with HALT^TM^ protease and phosphatase inhibitors added (Thermo Fisher Scientific, PI-78444). After transfer to 1.5ml tubes and sonication for lysis, cells were centrifuged at 15,000 rpm at 4°C for 5 minutes and lysates were then transferred to a new tube. Protein concentrations were determined by BCA assay as directed by the manufacturer (Pierce, P123227) and samples were prepared with Laemmli sample buffer (Bio-Rad, 161-0747), 100mM DTT (Sigma, D9779-5g) and heating at 95°C on a dry block heater for 5 minutes. Samples were separated by SDS-PAGE on a 4-12% Bis Tris gel (NuPage, NP0335BOX) and transferred onto a polyvinylidene fluoride membrane (Millipore, IPVH00010). Membranes were blocked with 5% bovine serum albumin (Sigma, A3912-100g) and incubated overnight at 4°C with primary antibodies [β GAPDH (Santa Cruz, sc-47724), cleaved PARP (CST, 5625S), cleaved caspase 3 (CST, 9661S), cleaved caspase 7 (CST, 9491S), HMOX-1 (CST, 43966S), GCLM (Abcam, ab126704), NQO1 (CST, 3187S), NRF2 (CST, 12721S), KEAP1 (CST, 8047S), PRDX1 (CST, 8499S), GSTO1 (Thermo, PA5-83383)] in 5% BSA in TBS-T (Bio-Rad, 170-6435). Following incubation with primary antibodies, membranes were washed three times for five minutes with TBS-T and incubated for one hour with 1:15,000 HRP-linked anti-rabbit or anti-mouse antibodies (Cell Signaling, 7074S, 7076S) or 1:15,000 DyLight anti-rabbit IgG 800 conjugate (Cell Signaling, 5366S). Membranes were then washed three times for five minutes with TBS-T and imaged with a Bio-Rad ChemiDoc MP imager or in a dark room on x-ray film (Pierce Biotech, PI34091). If HRP was used, either Pico or Fempto Chemiluminescent substrate (Pierce, PI34577/PI34095) was added prior to imaging. Redox western blots were performed by seeding 1.5 million cells in a 10cm plate and treating for two hours with each treatment condition the following day using established methods (39). Primary antibodies used: β sc-47724), cleaved PARP (CST, 5625S), cleaved caspase 3 (CST, 9661S), cleaved caspase 7 (CST, 9491S), HMOX-1 (CST, 43966S), GCLM (Abcam, ab126704), NQO1 (CST, 3187S), NRF2 (CST, 12721S), KEAP1 (CST, 8047S), PRDX1 (CST, 8499S), GSTO1 (Thermo, PA5-83383), PRDX1 (8499S).

### Annexin V assay

100,000 cells of each cell lines were seeded into 6-well plates. The day after seeding, cells were treated with 0.1% DMSO or 3µM LCS3 in triplicate. After 48 hours, cells were collected by trypsinization and prepared for flow cytometry according to manufacturer’s instructions (BD Bioscience, 556547).

### Transcript expression profiling (microarray)

H23 and H1650 cells were treated with DMSO for 3 or 6 hours in duplicate, or 3µM LCS3 for 3, 6, or 12 hours in duplicate, and profiled using the Affymetrix Human GeneChip 1.0 ST array platform as previously described (70). Raw data were normalized by robust multiarray analysis and a generalized linear model was applied via the rma and limma packages respectively (71, 72). Genes with a Benjamini-Hochberg adjusted p-value <0.05 in the LCS3 treated groups versus vehicle control were considered differentially expressed. Significantly upregulated genes in all treatment conditions were pooled and submitted to DAVID Bioinformatics Database v6.8 for functional analysis (73). We also used this upregulated transcript gene set to search the computational systems biology research tool Enrichr ENCODE TF ChIP-seq 2015 query function to predict the upstream regulators of LCS3-induced transcriptome changes (16, 17). The transcriptome dataset has been uploaded to NCBI Gene Expression Omnibus (GSE178311).

### Protein expression profiling (SILAC)

H23 cells were cultured either in normal media (light) or media with Lys and Arg substituted with ^13^C_6_ N_2_ Lys and ^13^C_6_ Arg (heavy) (Cambridge Isotopes). One replicate of each treatment condition was cultured in light media and one in heavy. Cells were maintained in label-containing media and passaged until 99% labeling efficiency was achieved. Cells were treated in duplicate (one heavy and one light) with 3µM LCS3 for 24 hours and then harvested by cell scraping. Samples were mixed 1:1, digested with trypsin, and prepared for mass spectrometry as previously described (15, 74). The SILAC proteome dataset has been provided as a supplementary file.

Normalized Z-scores were calculated for each identified protein replicate based on its ratio of treatment-to-vehicle (75). Z-score values were statistically combined by Stouffer’s Z method. Proteins with Z > 1.5 were considered differentially expressed. The upregulated proteins were entered as input into DAVID Bioinformatics Database v6.8 to determine upregulated biological processes by gene ontology (73).

### Antioxidant Response Element reporter assay

The ARE-GFP lentivirus and pathway control-GFP lentivirus (GenTarget, LVP981-B-PBS, Path-Ctr5-PBS) were transduced into H23 and H1650 cells with 8µg/ml polybrene (Sigma, H9268-10G) and selected with blasticidin (Life, R210-01). Two hundred thousand cells of H23 and H1650 pathway control-GFP and ARE-GFP cells were seeded into 6-well plates. One day after seeding, cells were treated with DMSO (0.1%), AI-1 (10µM), or LCS3 (1-3µM) in triplicate. After 72 hours, cells were washed with PBS, trypsinized and analyzed by flow cytometry.

### Detection of ROS

In 6-well plates, cells grown to 90% confluence were treated in triplicate with inhibitors. Thirty minutes into treatment, 10µM H_2_DCFDA (Enzo, 89152-832) was added as co-treatment to each well. After 1 additional hour, cells were washed twice with PBS, trypsinized, and stained with 1µg/ml propidium iodide (Thermo, P1304MP). H_2_DCFDA fluorescence (FL-1) was detected in propidium iodide-negative cells (FL-3) by a FACSCalibur Flow Cytometer (BD Biosciences).

### TPP

Five million H23 cells were seeded in two 15cm plates and harvested the next day by 2x PBS wash and gentle scraping using a cell lifter. Cells were pelleted at 1000rpm, resuspended in PBS, and lysed by three freeze-thaw cycles using a dry ice ethanol bath. Protein concentration was determined by BCA assay and diluted in PBS to a concentration of 1mg/ml. Lysates (1.2mL volume) were then treated with either 0.1% DMSO or 2µM LCS3 in duplicate and processed by the TPP protocol (12). Proteins were digested, reduced, alkylated and cleaned up by single-pot solid-phase-enhanced sample preparation (SP3) by established methods (76). In the multiplexed tandem mass tag (TMT) preparation (Thermo, A34808), in the 11^th^ channel (TMT11-131C) we included an untreated H23 lysate mix, which was not subjected to temperature challenge or ultracentrifugation, in an effort to enhance discovery of underrepresented peptides. Detected proteins in the TPP dataset were ranked by a non-linear regression data analysis (NPARC) using an open source R package (25). Proteins with an adjusted *p*-value < 0.01 were considered hits. This analysis produced a list of 77 protein candidates, which were then clustered into gene ontology biological processes by DAVID Bioinformatics Database v6.8 (73) for further analysis. The TPP dataset is provided as a supplementary file.

### Enzymatic assays

GSR activity was initially demonstrated by our lab group according to the Cayman Chemical Assay Kit protocol (703202) and then validated by an independent collaborator. Briefly, 20nM purified human GSR (Cedarlane, CLENZ202-2), was incubated with treatments in the presence of 800µM NADPH (Sigma, N1630-100MG) in 125µL assay buffer. To initiate the reaction, 75µL of 8mM oxidized GSSG (Sigma, G4376-1G) was added to each well and the absorbance was measured at 340nM for the duration of the experiment by a multi-mode imaging reader (BioTek, Cytation 3).

The laboratory of our collaborator, Dr. Katja Becker, independently performed GSR and TXNRD1 activity assays using established methods (33, 77). Human GSR was recombinantly produced in *E. coli* and purified using IPTG (Roth), Ni-NTA Agarose (Invitrogen, Karlsruhe) and Imidazole (Roth, Karlsruhe) (78) and native human TXNRD1 was isolated from human placenta (32). The GSR assay was performed by incubating purified recombinant GSR with NADPH (Biomol, Hamburg) and GSSG (Sigma-Aldrich, Steinheim) according to an established protocol (33). DTNB (Sigma-Aldrich, Steinheim) reduction assay was performed with isolated native human TXNRD1 using established methods (77). All measurements were conducted at least in two biological independent replicates and Lineweaver-Burk plots show the data of one representative experiment. IC_50_ values were calculated using the Quest Graph™ IC50 Calculator. AAT Bioquest, Inc, https://www.aatbio.com/tools/ic50-calculator.

### Docking studies

Protein preparation: 2GH5, a crystal structure of GSR complexed with a Fluoro-Analogue of the Menadione Derivative (1.70 Å resolution) and 2ZZ0, a crystal structure of TXNRD1 (2.80 Å resolution) were used for molecular docking studies. For protein structure preparation, all solvent molecules were deleted and the bond order for the protein was adjusted. Missing hydrogen atoms were added, and side chains were energy-minimized using the OPLS-2005 force field, as implemented by Maestro 2017v. The ligand binding region was defined by a 12 Å box centered on the crystallographic ligands of the crystal structures. Van der Waals scaling factors were not applied; the default settings were used for all other adjustable parameters. Ligand preparation: LCS3 was built using MOE version 2017. Hydrogen atoms were added after these structures were “washed” (a procedure including salt disconnection, removal of minor components, deprotonation of strong acids, and protonation of strong bases). The following energy minimization was performed with the MMFF94x force field, as implemented by the MOE, and the optimized structures were exported into the Maestro suite in SD file format. Molecular Docking: docking experiments were performed using Glide included in the Schrodinger Package, Maestro interface. For docking, standard-precision (SP) docking method was adopted to generate the minimized pose, and the Glide scoring function (Glide Score) was used to select the final poses for ligands.

### Measurement of GSH, GSSG, NAPDH, and NADP+ levels

For each cell line, a total of 5,000 cells/well were seeded into 96-well plates. Twenty-four hours later, media was replaced with media containing DMSO or specified concentrations of LCS3. After 6 hours of treatment, cells were washed with 150µL of PBS. GSH, GSSG, NADPH, and NADP+ levels were then quantified using either the GSH/GSSG-Glo assay (Promega) or the NADP/NADPH-Glo assay (Promega) according to manufacturer instructions.

### Genome-wide loss-of-function CRISPR-Cas9 knockout screen

Genome-wide screens were performed with the Toronto Knockout version 3 (TKOv3) library (47). Lentivirus was generated from the TKOv3 library in low passage (<10) 293FT cells (Thermo Fisher) using Lipofectamine 3000 (Thermo Fisher). Approximately 400 million target cells (H358s) were then infected with the TKOv3 library virus at a multiplicity of infection (MOI) of 0.5, in order to achieve an average 200-fold representation of the sgRNAs in each condition and replicate of the screen. Cells were selected on puromycin for 5 days and then 14 million cells were seeded into media containing DMSO, LCS3 4µM, or LCS38 µM in triplicate. Cells were passaged every 3 days and after 14 population doublings in the DMSO condition, 14 million cells from each condition were harvested for genomic DNA extraction. sgRNA inserts were amplified with NEBNext High-Fidelity 2X PCR Master Mix (New England BioLabs). Samples were then pooled in equimolar concentrations, gel purified, and sequenced on a NextSeq 500 kit (Illumina).

Sequencing reads were trimmed and aligned to the reference library using Cutadapt and Bowtie, respectively. This alignment returned a raw read count table, and the sgRNAs with less than 30 raw read counts were excluded from further analysis. The raw read counts were then analyzed with MAGeCK-MLE in order to obtain a beta-score for each gene (79). To identify gene knockouts that conferred resistance to LCS3, the beta-scores for the genes in the LCS3-treated conditions (4 or 8µM) were subtracted from the beta-scores in the DMSO-treated condition. The CRISPR screen dataset is provided as a supplementary file.

### siRNA knockdown

400,000 cells were first seeded into a 6-well plate. The next day, cells were transfected with siRNA by combining 10µl of 20µM siRNA and 5µL Dharmafect (Dharmacon, T-2001-03) in 800µL OptiMEM (Gibco, 31985-062) and adding to a 6-well plate in RPMI media to a total volume of 4mL, for a final siRNA concentration of 50nM. The siRNAs used were non-targeting pool siCtrl (Dharmacon, D-001810-10-05), siGSR (Dharmacon, E-009647-00-0005), siTXNRD1 (Dharmacon, L-008236-00-0005), and siKIF11 (Dharmacon, L-003317-00-0005).

### In-gel fluorescence assay

In 200µL PCR tubes, 0.1% DMSO, LCS3, or C1-27 was added to 0.5µg of recombinant GSTO1 (Sigma, GS75-100ug) in PBS in 12uL total volume and incubated in a thermocycler at 37°C for 20 mins in a competitive CMFDA binding assay described previously (28). Six µL of 3µM CMFDA (Cedarlane, G783003) was then added to each sample and incubated for an additional 20 mins. Samples were heated in a thermocycler at 95°C for five minutes and mixed with Laemmli sample buffer and DTT and loaded into a 4-12% Bis-Tris gel. Samples were separated on the gel and entire gel was imaged on a Bio-Rad ChemiDoc MP on the fluorescein image setting. The gel was then transferred to PVDF and blotted for GSTO1 as described in western Blot methods section.

## Acknowledgements

This work was funded by the BC Cancer Foundation, the Cancer Research Society, Canadian Institutes of Health Research (CIHR; PJT-148725) and the Terry Fox Research Institute (New Investigator Award) to W.W.L.

## Author contributions

Conceptualization, F.D.J, H.V, G.B.M, P.H.S., J.F., A.L. and W.W.L. Methodology, F.D.J, J.F., A.L., C.B., K.B., S.J.D., R.M., S.E.S.M, G.S.W.C., G.B.M, H.D., U.G., X.Z. and W.W.L.; Validation, C.B. and K.B.; Investigation, F.D.J., J.F., A.L., C.B. D.F., D.L., S.J., J.L., T.S., R.S., G.C.F., G.R. and W.W.L.; Formal Analysis, F.D.J., J.F., R.M., A.L. and W.W.L; Visualization, Y.I., A.N., D.F., M.G., J.L., R.M. and W.W.L.; Writing – Original Draft, W.W.L., F.D.J.; Writing – Review & Editing, W.W.L., J.F, H.V, C.B., R.S., H.D., S.J.D., G.B.M., S.E.S.M.; Funding Acquisition, W.W.L.; Resources, H.V and W.W.L; Supervision, H.V, G.B.M., and W.W.L.

## Supplemental Materials

**Supplementary Figure 1. a,** Incucyte images taken at 72 hours of treatment with LCS3 or GPX4 inhibitor RSL3 (positive cell death control). Cell lines expressing nuclei marker H2B-mCherry (red) and treated with dead cell marker Sytox Green (green). LCS3-resistance cell lines in bold font. **b,** Calculation of D_o_ and D_R_ for 4µM LCS3 in all STACK cell lines. **c,** IC_50s_ values for inhibition of growth of the NCI-60 cell line panel by LCS3 was determined by the pipeline established by the Developmental Therapeutics Program at the NCI.

**Supplementary Figure 2. a,** Volcano plots of microarray data showing significant differentially expressed genes in H23 and H1650 cells treated with 3µM LCS3 for 3, 6, or 12 hours relative to control. **b,** Z-score plot of SILAC data showing significant differentially expressed genes in H23 treated with 3µM LCS3 for 24 hours. **c,** Schematic of NRF2 pathway.

**Supplementary Figure 3. a,** In-gel fluorescence assay (top section) to determine GSTO1 active site occupancy by LCS3 and positive control GSTO1 inhibitor C1-27. Lower section is a western blot showing total GSTO1 in each lane. **b,** Incucyte growth experiment of both LCS3-sensitive and LCS3-resistant cell lines treated with either LCS3 or GSTO1 inhibitor C1-27. **c,** Proliferation screen of 24 lung cell lines and 27 LCS3 analogues. Nitro subgroup-containing compounds are indicated by a gray box. **d,** *In vitro* enzymatic GSR activity assay of LCS3 and an inactive analogue which lacks the nitro functional group.

**Supplementary Figure 4 a,** Heatmaps showing combination interactions between inhibitors of the glutathione and thioredoxin pathways. **b,** Incucyte growth experiment demonstrating interaction between GSR depletion by siRNA and either LCS3 or TXNRD1 inhibitor auranofin. Data are shown as mean +/- SEM. **c,** IncuCcyte growth experiment demonstrating the lack of combination synergy between BSO and TXNRD1 depletion by siRNA.

**Supplementary Figure 4 a,** Western blot of sgNQO1 cell lines and control cell line sgChr19-SAFE treated with LCS3 or DMSO for 6 hours. **b,** Metabolite assay to determine the relative concentrations of NADPH and NADP^+^ in H358. Error bars are +/- SD. **c,** Incucyte growth experiment demonstrating that KEAP1 deletion does not impact LCS3 sensitivity in H358 cells. Note that sgChr19-SAFE control dose-response data are same as those shown in Fig. 6b Incucyte with sgNQO1 cell lines. Data are shown as mean +/- SEM. **d,** Western blot showing NRF2 pathway activation of H358 cells with KEAP1 deletion at the basal state.

**Supplementary Table 1.** LCS3 derivative structures used in SAR investigation (Supplementary Fig. 3c).

**Supplementary Table 2.** SILAC dataset of experiment shown in Fig. 2a.

**Supplementary Table 3.** TPP dataset and NPARC analysis

**Supplementary Table 4.** Genome-wide CRISPR-Cas9 knockout screen dataset.

