## Supplemental Figures 1-5 for "A novel small molecule that induces cytotoxicity in lung cancer cells inhibits disulfide reductases GSR and TXNRD1"

## 1.a

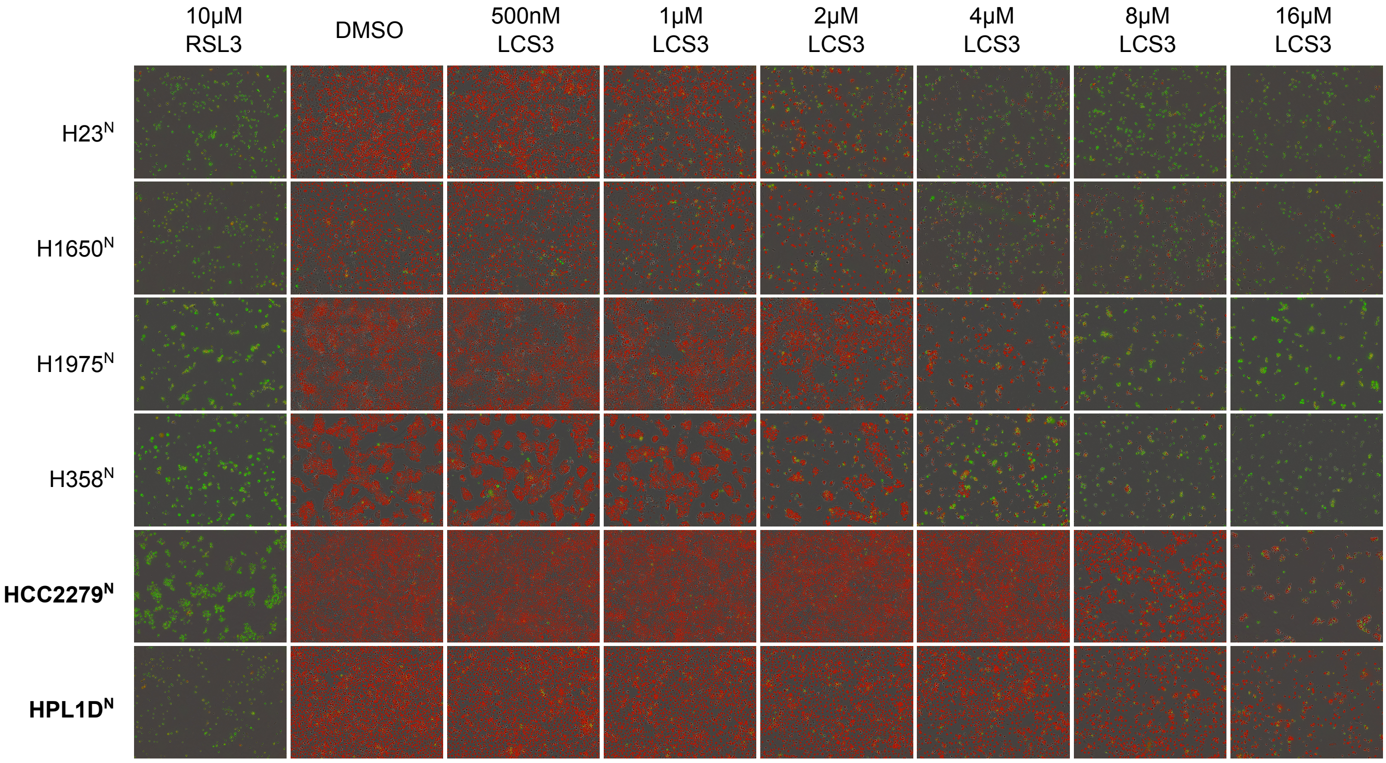


## 1.b

| | LCS3 (4μM) | | | | | |
|---|---|---|---|---|---|---|
| | $H23^{N}$ | $H1650^{N}$ | $H1975^{N}$ | $H358^{N}$ | **$HPL1D^{N}$** | **$HCC2279^{N}$** |
| $D_o$ (hrs) | 9.1 | 6.0 | 17.1 | 24.9 | 27.1 | -- |
| $D_R$ (% dead $hr^{-1}$) | 2.2 | 1.2 | 1.5 | 7.4 | 0.6 | -- |

## 1.c

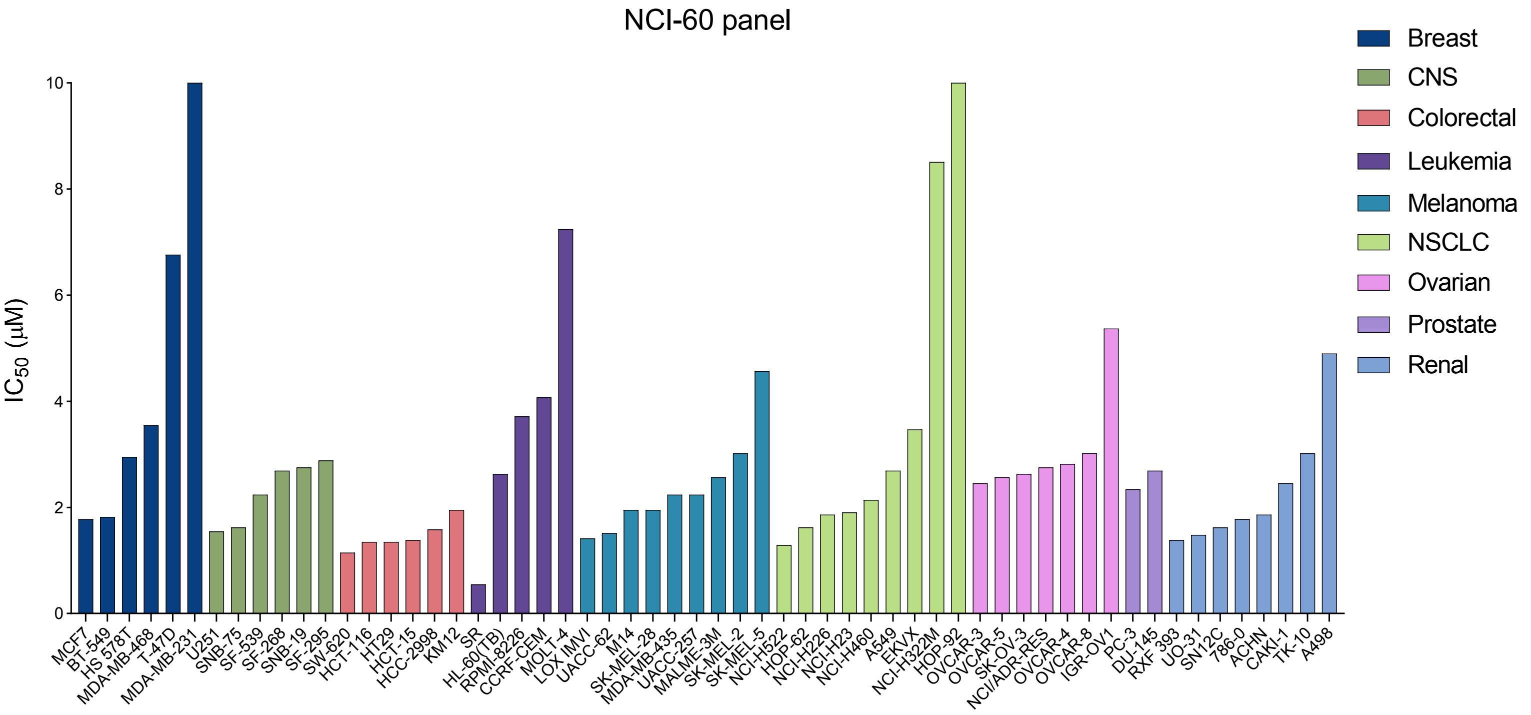

### Supplemental figures

#### Microarray

## 2.a

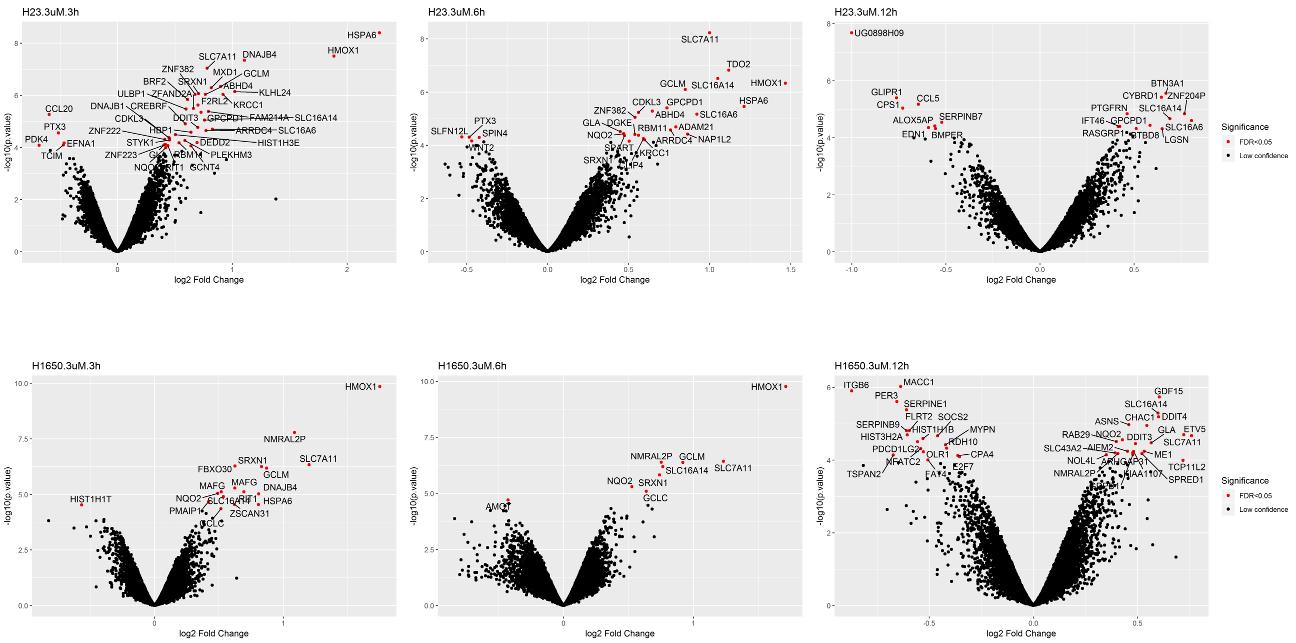


## 2.b

#### SILAC

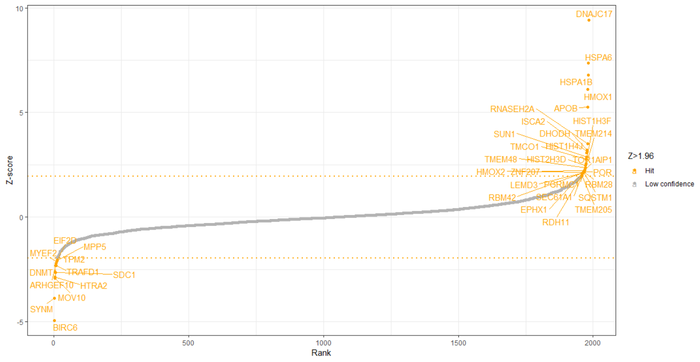


## 2.c

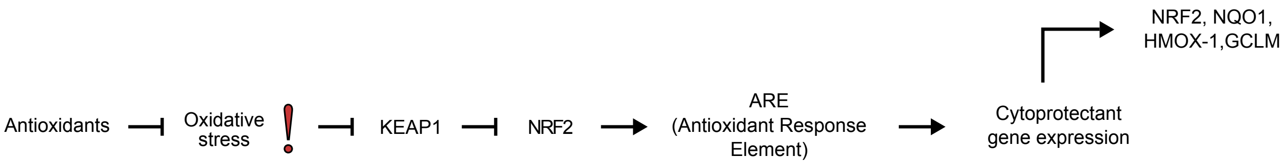

### Supplemental figures

## 3.a

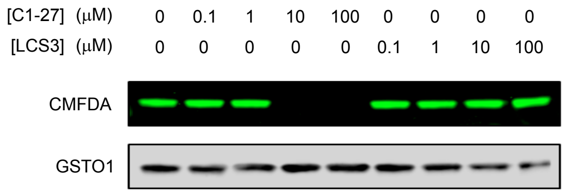


## 3.b

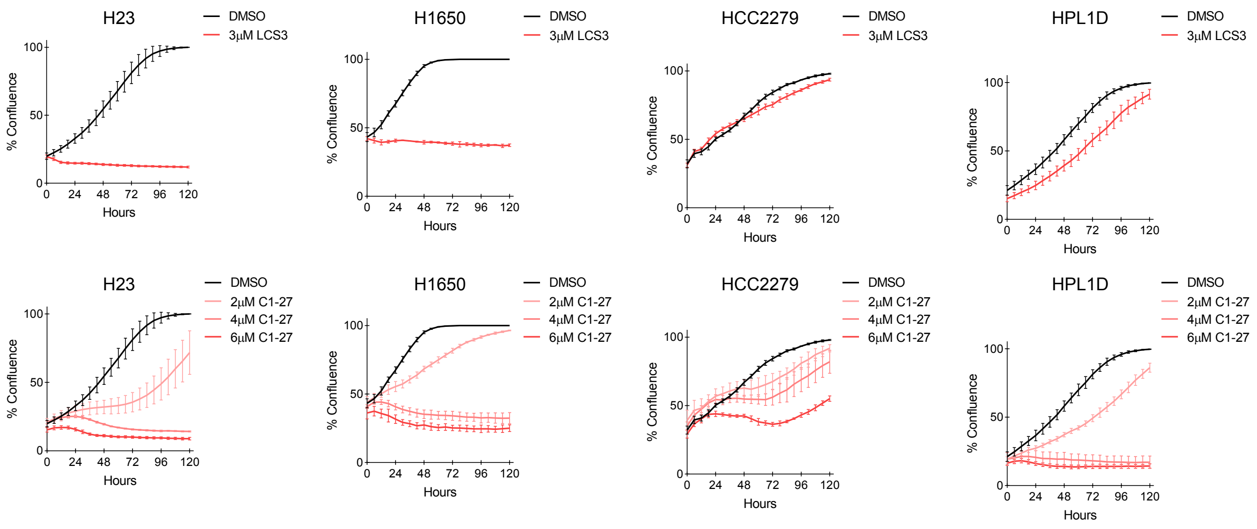


## 3.c

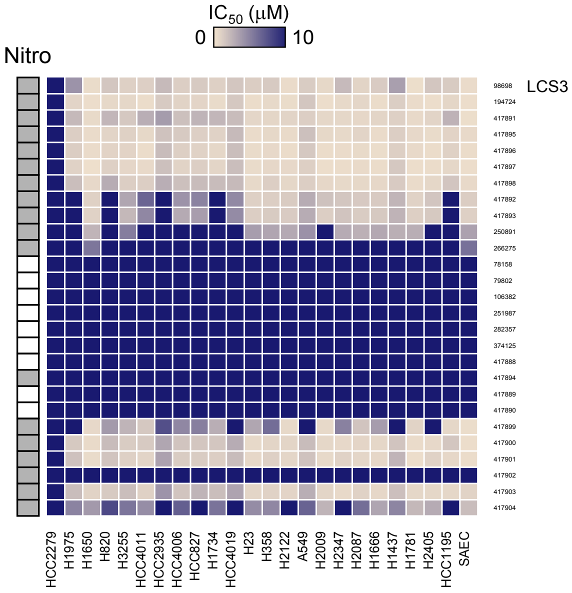


## 3.d

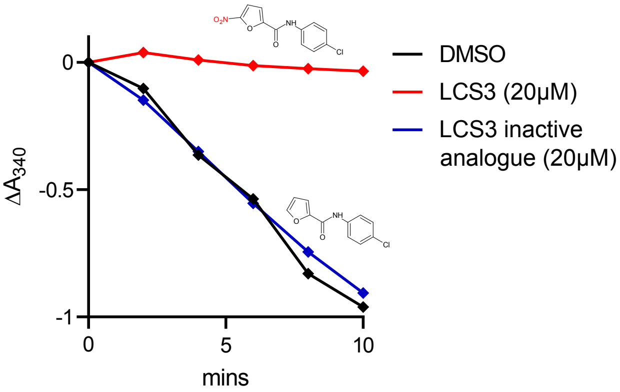

### Supplemental figures

4.a

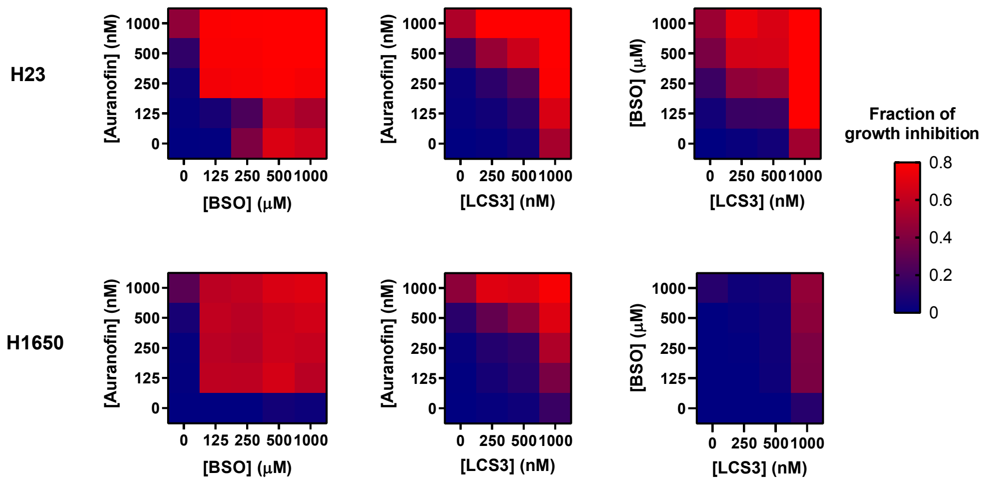


4.b

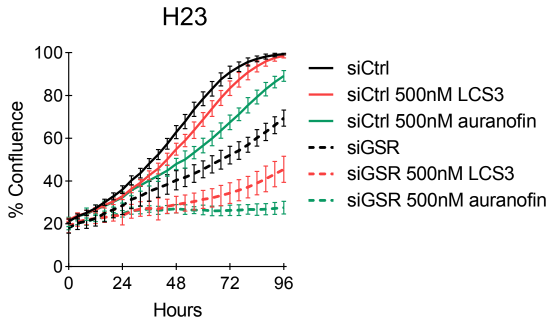


4.c

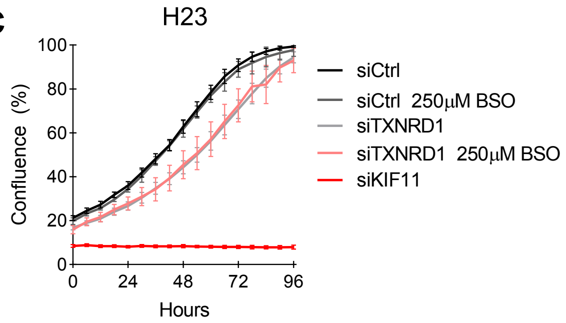

### Supplemental figures

5.a

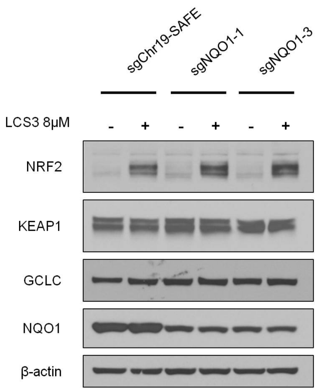


5.b

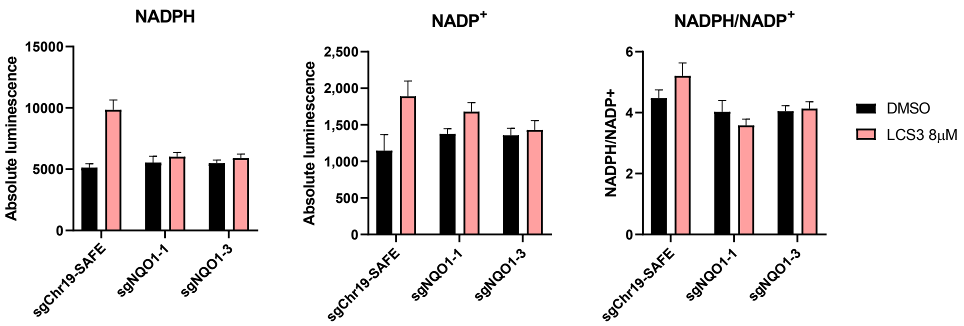


5.c

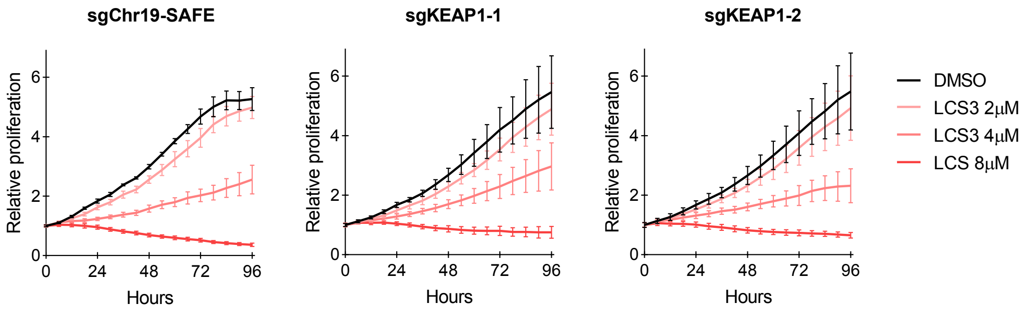


5.d

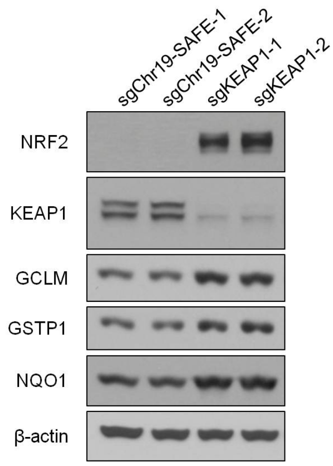
