## Supplemental Table 1 for "A novel small molecule that induces cytotoxicity in lung cancer cells inhibits disulfide reductases GSR and TXNRD1"

### LCS3 derivative structures

| | | |
|---|---|---|
| 98698 (LCS3)<br>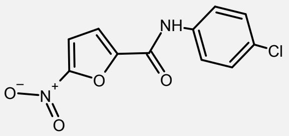 | 250891<br>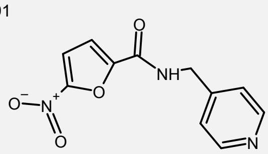 | 417894<br>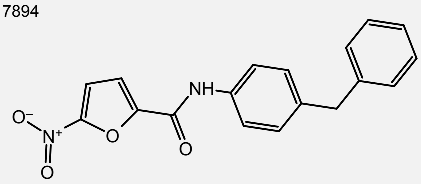 |
| 194724<br>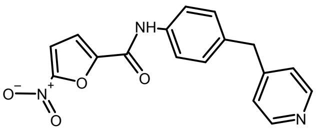 | 266275<br>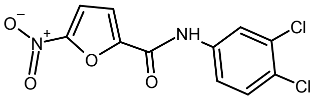 | 417889<br>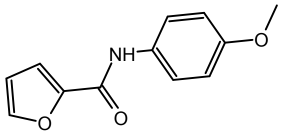 |
| 417891<br>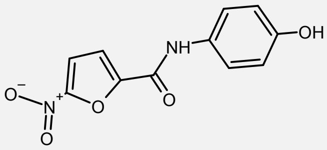 | 78158<br>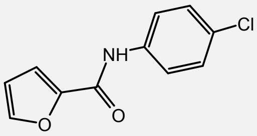 | 417890<br>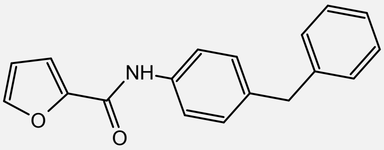 |
| 417895<br>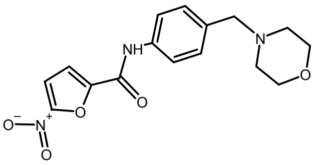 | 79802<br>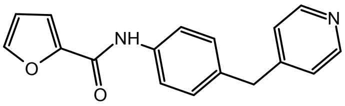 | 417899<br>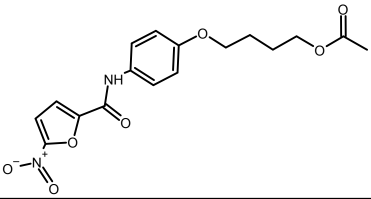 |
| 417896<br>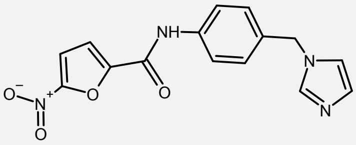 | 106382<br>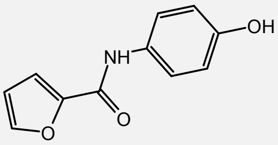 | 417900<br>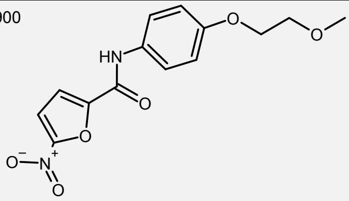 |
| 417897<br>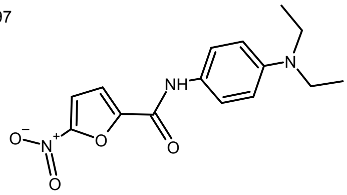 | 251987<br>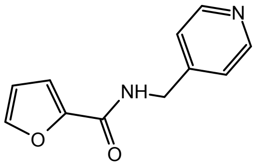 | 417901<br>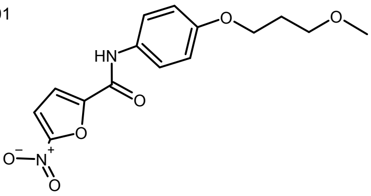 |
| 417898<br>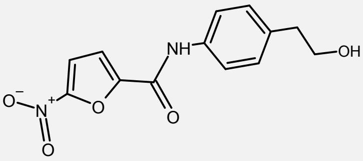 | 282357<br>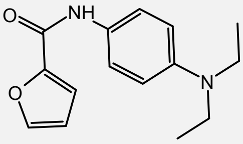 | 417902<br>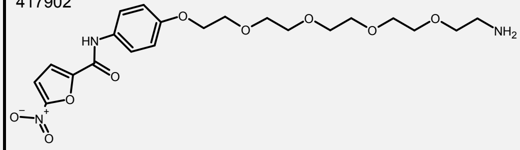 |
| 417892<br>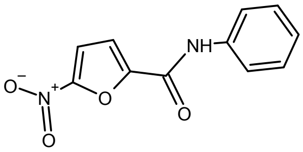 | 374125<br>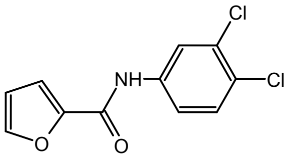 | 417903<br>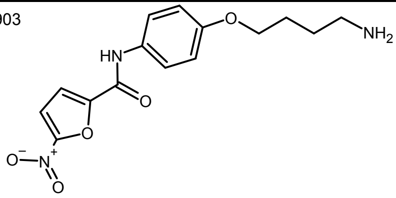 |
| 417893<br>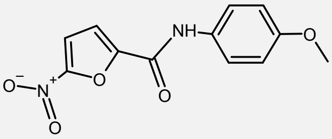 | 417888<br>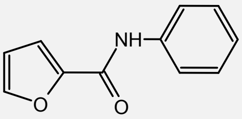 | 417904<br>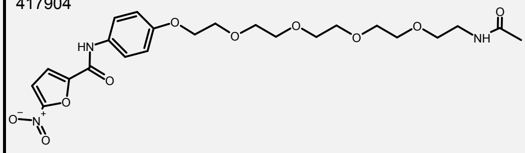 |
